# *In vivo* metabolomics identifies CD38 as an emergent vulnerability in *LKB1*-mutant lung cancer

**DOI:** 10.1101/2023.04.18.537350

**Authors:** Jiehui Deng, David H. Peng, David Fenyo, Hao Yuan, Alfonso Lopez, Daniel S. Levin, Mary Meynardie, Mari Quinteros, Michela Ranieri, Soumyadip Sahu, Sally C. M. Lau, Elaine Shum, Vamsidhar Velcheti, Salman R. Punekar, Natasha Rekhtman, Catríona M. Dowling, Vajira Weerasekara, Yun Xue, Hongbin Ji, Yik Siu, Drew Jones, Aaron N. Hata, Takeshi Shimamura, John T. Poirier, Charles M Rudin, Takamitsu Hattori, Shohei Koide, Thales Papagiannakopoulos, Benjamin G. Neel, Nabeel Bardeesy, Kwok-kin Wong

## Abstract

LKB1/STK11 is a serine/threonine kinase that plays a major role in controlling cell metabolism, resulting in potential therapeutic vulnerabilities in LKB1-mutant cancers. Here, we identify the NAD^+^ degrading ectoenzyme, CD38, as a new target in LKB1-mutant NSCLC. Metabolic profiling of genetically engineered mouse models (GEMMs) revealed that LKB1 mutant lung cancers have a striking increase in ADP-ribose, a breakdown product of the critical redox co-factor, NAD^+^. Surprisingly, compared with other genetic subsets, murine and human LKB1-mutant NSCLC show marked overexpression of the NAD+-catabolizing ectoenzyme, CD38 on the surface of tumor cells. Loss of LKB1 or inactivation of Salt-Inducible Kinases (SIKs)—key downstream effectors of LKB1— induces CD38 transcription induction via a CREB binding site in the CD38 promoter. Treatment with the FDA-approved anti-CD38 antibody, daratumumab, inhibited growth of LKB1-mutant NSCLC xenografts. Together, these results reveal CD38 as a promising therapeutic target in patients with LKB1 mutant lung cancer.

**SIGNIFICANCE:** Loss-of-function mutations in the *LKB1* tumor suppressor of lung adenocarcinoma patients and are associated with resistance to current treatments. Our study identified CD38 as a potential therapeutic target that is highly overexpressed in this specific subtype of cancer, associated with a shift in NAD homeostasis.

## INTRODUCTION

The *STK11* tumor suppressor gene encodes a serine-threonine kinase, *LKB1*, with pleiotropic functions in cell metabolism, polarity, and growth control. Germline loss-of-function *STK11* mutations result in Peutz-Jeghers Syndrome (PJS), characterized by gastrointestinal polyposis and increased risk of various malignancies (1). Moreover, somatic mutations and deletions of *LKB1* are observed in many sporadic tumor types, including lung cancer (2–4). LKB1 inactivation is particularly common in lung adenocarcinoma, with ∼15-20% of cases exhibiting genomic inactivation of *LKB1* (4–7). This subset of non-small cell lung cancer (NSCLC) is associated with poor outcomes and reduced sensitivity to existing conventional chemotherapies and immunotherapies compared with *LKB1* WT tumors (8,9). LKB1 phosphorylates and activates the 14 members of the AMPK-activated protein kinase (AMPK) subfamily, many of which are involved in metabolic regulation (10). Functional studies indicate that, among this group, the Salt-Inducible kinases (especially SIK1 and SIK3) are important for tumor suppression by LKB1, in keeping with the emerging importance of SIKs in growth regulation (11–13). Despite extensive preclinical and clinical studies of LKB1-mutant NSCLC, treatment of this subset of tumors remains a major unmet need in oncology.

CD38 is a type II transmembrane receptor expressed in a variety of tissues, including lymphatic and myeloid lineages, where it regulates cell proliferation and immune response, as well as in bronchial epithelial cells, pancreatic islet cells, and others (14,15). This multifunctional ectoenzyme serves as an NAD^+^ glycohydrolase and cyclic ADP-ribose synthase, and generates ADP-ribose—a metabolite also produced by other NAD^+^ consuming enzymes, including sirtuins (SIRT1-5) and poly(ADP-ribose) polymerases (PARPs) (16,17). NAD^+^ is also an essential coenzyme that coordinates electron transfer critical for diverse metabolic redox reactions (18,19). CD38 regulates global NAD^+^ availability through its ability to continuously degrade NAD^+^ (20). Multiple myeloma exhibits prominent overexpression of CD38, and accordingly, anti-CD38 antibodies show substantial clinical efficacy against this malignancy (21,22). In this study, we performed a systemic analysis of metabolic changes in LKB1 mutant lung cancers and identified CD38 as a therapeutic target in this setting. Furthermore, we found that the SIK family kinases were crucial for the suppression of CD38 expression at transcriptional level via CRTC/EP300/CREB complex. These findings suggest the potential of employing currently available therapeutic antibodies against CD38 for the treatment of LKB1 mutant lung cancer, which currently lack of treatment strategies.

## RESULTS

### LKB1 loss-of-function shifts metabolism of NAD homeostasis

The impact of LKB1 deficiency on cell metabolism has been studied extensively *in vitro*, but little is known about the metabolic profiles of *STK11*-mutant tumors *in vivo*. To address this issue, we performed targeted metabolomic profiling of NSCLC allografts generated using cell lines from *Kras^G12D^*; *Lkb1*^-/-^ (KL)(23) and *Kras^G12D^Trp53*^-/-^ (KP)(24) genetically engineered mouse models (GEMMs) implanted in syngeneic immunocompetent mice. Liquid chromatography/mass-spectrometry (LC/MS) revealed largely overlapping profiles between genotypes for the 147 metabolites tested. Notable exceptions were three metabolites showing highly statistically significant increases in expression in KL tumors and 11 showing significant elevations in KP tumors. Adenosine diphosphate ribose (ADP ribose) and glutamate had the highest fold-changes among the upregulated metabolites in KL tumors (**Fig. 1A** and **Table 1**). ADP ribose is generated from catabolism of the redox co-factor, NAD^+^, by Sirtuins, PARP family member, and CD38 (25–27). Notably, RNAseq demonstrated increased CD38 expression in KL tumors compared with KP tumors, whereas PARP, Sirtuin, and NAD+ biosynthesis and salvage pathway enzyme transcript levels were comparable in the two genotypes (**Supplementary Fig. S1A-S1C**). Flow cytometric analysis of tumor tissue demonstrated increased cell surface CD38 expression on neoplastic (EpCAM+) cells from the KL model, with both a higher proportion of CD38+ cells and increased total CD38 expression compared with the KP model (**Fig. 1B, 1C**). Furthermore, examination of 92 KRAS-mutant human NSCLCs from TCGA revealed increased CD38 mRNA levels in the subset with *LKB1* co-mutations (N=32) compared with *LKB1* wild type (N=45) tumors (**Fig. 1D, Supplementary Table S1**). In line with findings from the GEMMs, CD38 was prominently upregulated in human KL tumors (**Supplementary Fig. S1B**), whereas other NAD consuming and biosynthesis/salvage enzymes did not show significant differences between genotypes (**Supplementary Fig. S1A-S1C, Supplementary Table S2**). Importantly, immunohistochemical analysis of NSCLC patient samples demonstrated a significant increase in CD38 staining score in the *LKB1* mutant tumors (*p*=0.0002), corroborating our findings from the human cell line CCLE DepMap database (**Fig. 1E, 1F, Supplementary Table S3, Supplementary Fig. S1B-S1C**).

**Figure 1.**
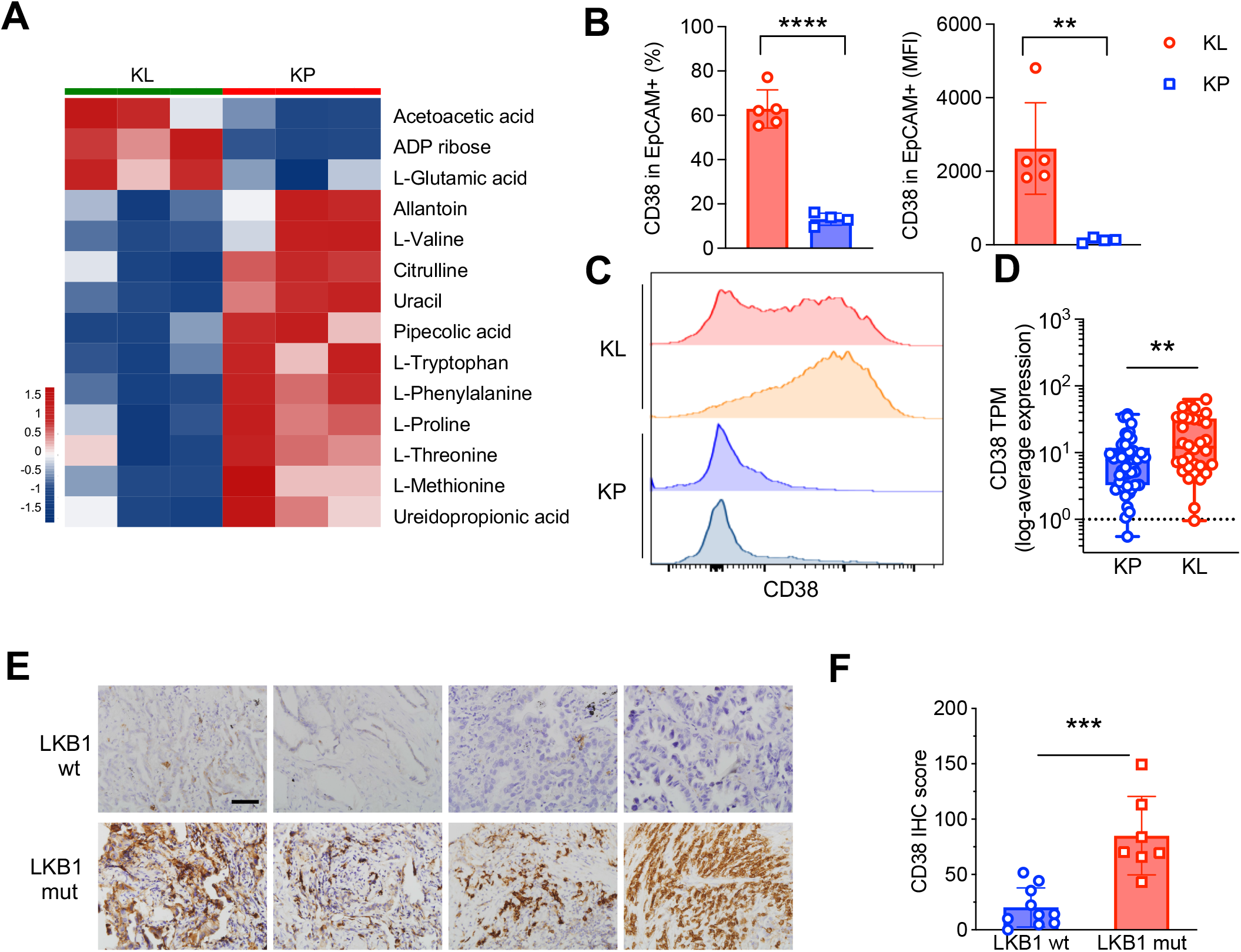
LKB1 deficient NSCLC show shifts in NAD^+^ homeostasis and upregulation of the CD38 NADase. **A**, Metabolites that show significant changes in KL mouse lung tumors compared to KP tumors. Values are shown as z score of fold-change. **B**, **C,** Flow cytometry analysis showing quantification of CD38 expression in neoplastic (EPCAM+) cells in *Kras*/*Lkb1* (KL) and *Kras*/*Trp53* (KP) tumors. **B**, (left panel) percent positive tumor cells and (right panel) median fluorescent intensity (MFI). Student’s *t* test, two-tailed. \*\**P*<0.01, \*\*\*\**P*<0.0001. **C**, Representative FACS blots of CD38 expression. **D**, *CD38* mRNA levels in human KP and KL lung cancer patient specimens, from TCGA dataset. Unpaired *t* test with Welch’s correction, two-tailed. \*\**P*<0.01. **E,F,** Lung cancer patient specimens with LKB1 wt and mutant status were stained with anti-CD38 antibodies. **E**, Representative IHC images. Scale bar, 100 μm. **F**, Quantification of CD38 IHC score. Student’s unpaired *t* test, two tailed. \*\*\**P*<0.001.

### CD38 is activated in LKB1 mutant lung cancer cells

We used a series of human KRAS mutant NSCLC cell lines with or without LKB1 alteration to further investigate the relationship between LKB1 mutations and CD38 expression. CD38 surface levels were markedly (∼5-50-fold) higher in 3 of 4 lines with genomic inactivation of *LKB1* (HCC44 [frameshift insertion], A549 [nonsense mutation], and H1944 [missense mutation, K62N]) compared with *LKB1* WT lines (H441, H358, Calu1, and H1792) (**Fig. 2A-2C, Supplementary Fig. S2A, Supplementary Table S2**). Flow cytometric analysis of PDX models corroborated these findings *in vivo*, showing high levels of cell surface expression of CD38 in *LKB1* mutant Lx337 (*KRAS*^G12C^*STK11*^Del^) tumors relative to *LKB1* wild type Lx210 (*KRAS*^G12V^*STK11*^wt^) tumors (**Fig. 2D, 2E, Supplementary Fig. S2B**). Importantly, restoring WT LKB1 expression in A549 cells suppressed CD38 expression, whereas a kinase-dead mutant of LKB1 (LKB1-KD) failed to do so (**Fig. 2F, 2G**). Thus, LKB1 is a negative regulator of CD38.

**Figure 2.**
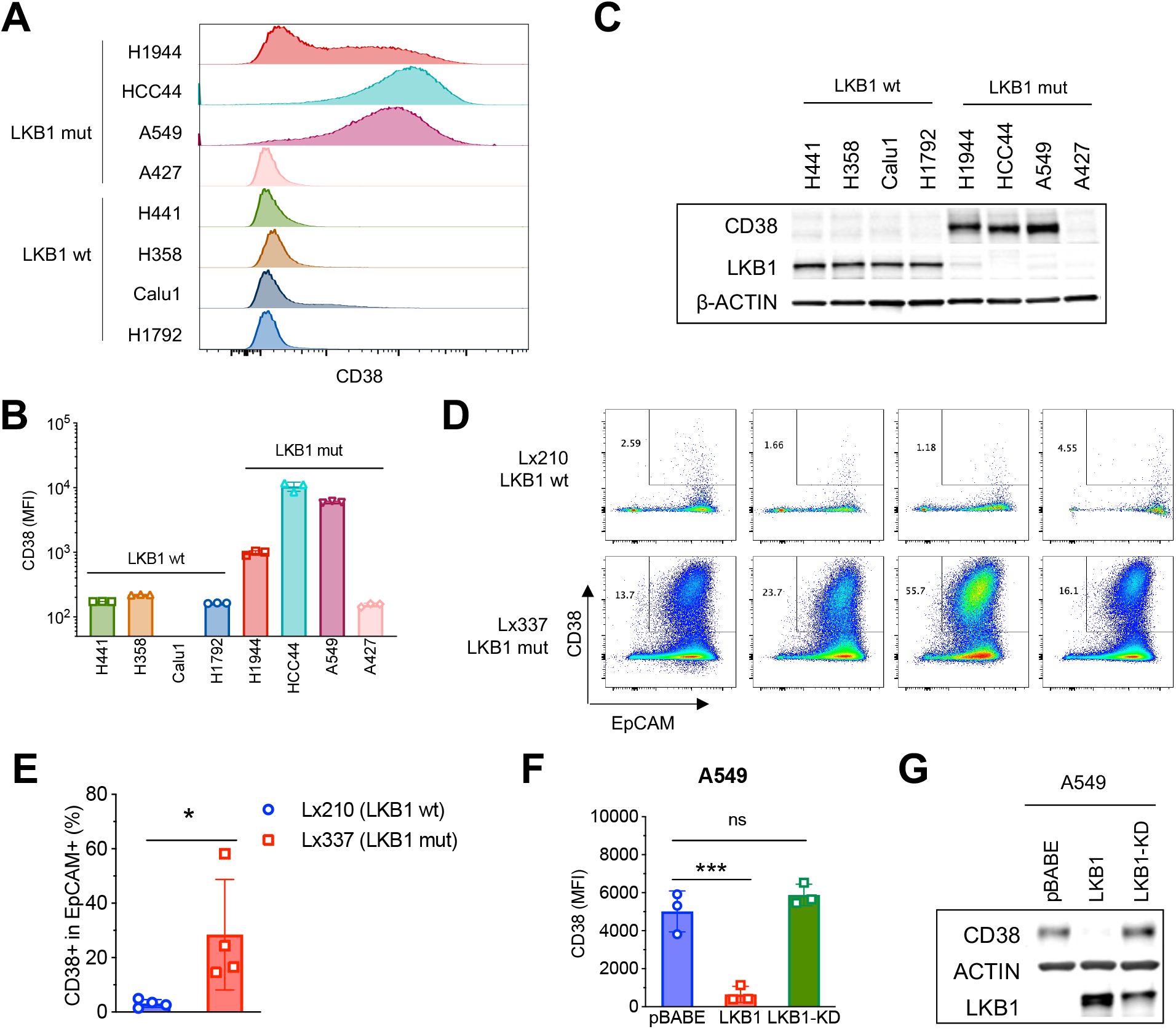
LKB1 is a negative regulator of CD38 expression in lung cancer cells. **A**,**B**, A series of *KRAS* mutant NSCLC cell lines with or without *LKB1* mutation were analyzed for CD38 expression via FACS. **A**, Representative FACS plots. **B**, Quantification. n=3 biological replicates for each cell line. **C**, Western blot analysis of CD38 protein levels in *KRAS* mutant lung cancer cell lines with or without *LKB1* mutation. **D**, **E,** *LKB1* mutant (Lx337) and *LKB1* wildtype (Lx210) lung cancer PDXs were tested for CD38 expression by FACS analysis. **D**, Representative FACS plots. **E**, Quantification of percent CD38+ tumor cells in the PDXs. n=4 tumors per group. Student’s *t* test, two tailed. \**P*<0.05. **F, G,** A549 NSCLC cells were transduced with empty vector (pBABE) or ectopically expressing *LKB1* wildtype (LKB1) or kinase-dead mutant (LKB1-KD). **F**, Cells were tested for CD38 expression by FACS analysis. n=3 each group. Each dot represents one independent experiment. One-way ANOVA Dunnett’s multiple comparisons test compared with pBABE group. \*\*\**P*<0.001. **G**, Western blot analysis of CD38 protein levels.

### Loss of SIK family kinase signaling is responsible for CD38 activation in LKB1 mutant tumors

We next examined the mechanism by which LKB1 regulates CD38, focusing on the SIK family kinases given their central role in LKB1-mediated tumor suppression (12,13). LKB1 activates SIKs by phosphorylating a threonine residue in the T-loop of their kinase domains (28). Accordingly, global phosphoproteomic analysis of human NSCLCs showed that *LKB1* mutant tumors have significantly lower levels of activating SIK phosphorylation (gauged by SIK3-T221 phosphorylation) than other *LKB1* wildtype lung tumors in the Clinical Proteomic Tumor Analysis Consortium (CPTAC) dataset (29) (**Fig. 3A, Supplementary Table S4**; the equivalent phospho-sites on the SIK1 and SIK2 T-loops were not detectable in this analysis). Consistent with decreased SIK kinase activity and the established feedback loop between SIK inactivation and transcriptional upregulation of SIK genes, we found that SIK1 and SIK2 mRNA levels were increased in human KL tumors compared with KP tumors (**Supplementary Fig. S3A**). Expression of the entire set of 14 LKB1 downstream AMPK/SIK kinases (10,28) was shown for reference and demonstrated that select other family members are also differentially expressed (e.g., *Ampk*/*Prkaa1*). Paralleling the human data, *Sik1* and *Sik2* were elevated in KL murine lung tumors and cell lines, along with *Sik3* and *Ampk*/*Prkaa1* (**Supplementary Fig. S3B, S3C**). Moreover, levels of CD38 correlated with SIK1 and SIK3, but not SIK2, in *LKB1* wild type lung cancer patients, but not in *LKB1* mutant patients (Supplementary Fig. S3D-S3F). This indicates SIK1 and SIK3 might be more important for mediating CD38 expression downstream of LKB1, but not SIK2.

**Figure 3.**
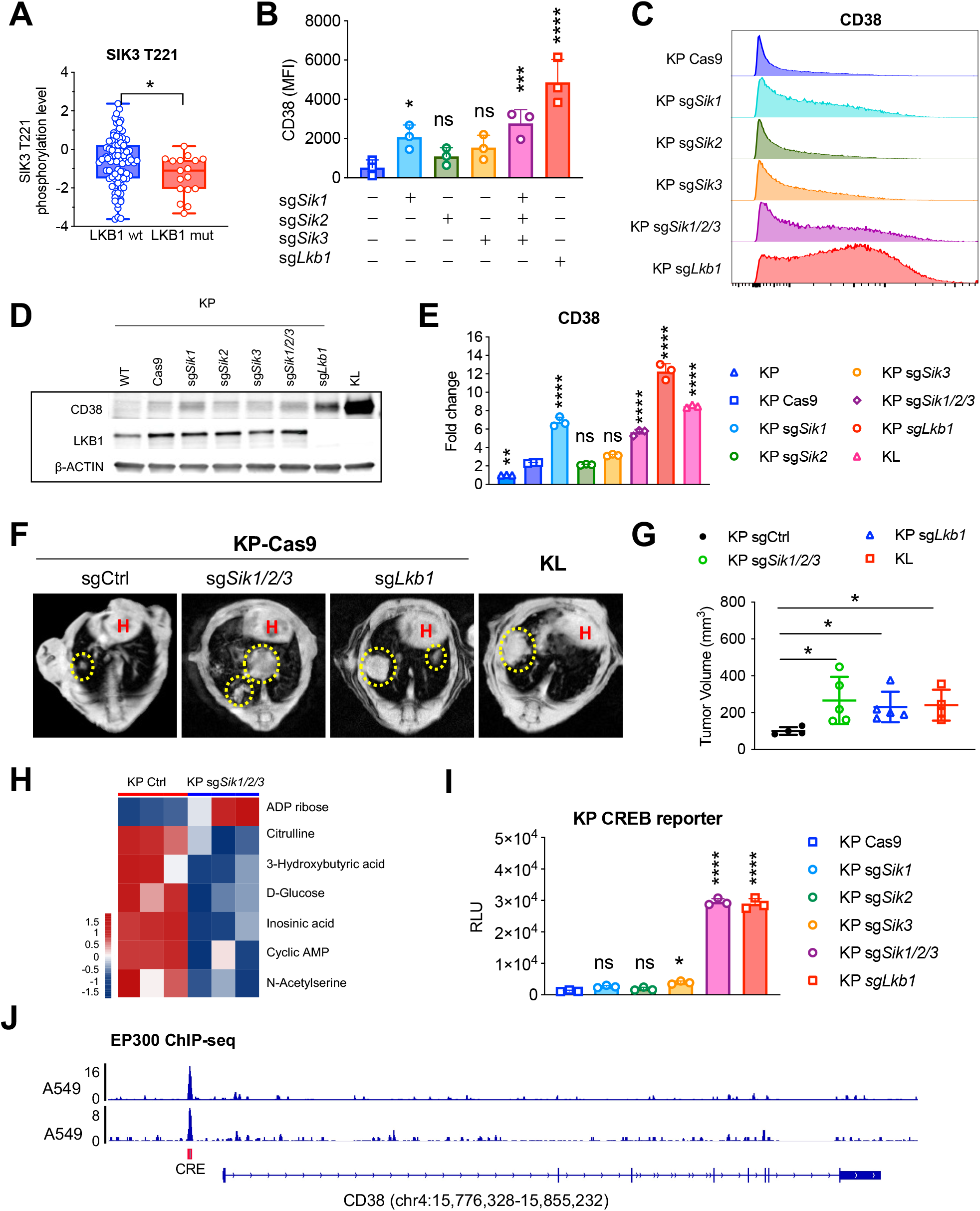
LKB1-SIK signaling suppresses CD38 transcriptionally. **A**, Relative phosphorylation levels of SIK3 T221 in human lungs cancer specimens with the indicated LKB1 genotypes (from the CPTAC dataset). Student’s unpaired *t* test, two tails. \**P*<0.05. **B,** FACS analysis showing CD38 expression on KP lung tumors with sgRNA mediated targeting of *Lkb1* or *Sik* family members versus control. \**P*<0.05, \*\*\**P*<0.001, \*\*\*\**P*<0.0001. **C**, Representative FACS analysis of CD38 levels in KP lung cancer cells shown in (**B**). **D**, Western blot of CD38 levels in KP lines without *Lkb1* or *Sik* family kinases expression, or KL lines without *Lkb1* expression. **E**, Relative CD38 mRNA levels in the lines shown in panel **B**. One-way ANOVA, Dunnett’s multiple comparisons test compared with KP Cas9. \*\**P*<0.01, \*\*\*\**P*<0.0001. **F**, Representative MRI scan of lung tumor nodules (circled) with the indicated genotypes. H, heart. **G**, Quantification of tumor volume changes 6 weeks after tumor implantation. Unpaired student’s *t* test in comparison with KP sgCtrl group. \**P*<0.05. **H**, Metabolite changes with statistical significance comparing mouse KP lung tumor nodules with or without *Sik1/2/3* loss. **I**, CREB reporter assay in the indicated cells. n=3 each cell line. Ordinary one-way ANOVA Dunnett’s multiple comparison test, each group compared with the KP Cas9 group. n=3 each group. \**P*<0.05, \*\*\*\**P*<0.0001. **J**, EP300 ChIP-seq track of CD38 promoter using the A549 cell line from the ENCODE database.

In line with the results of the LKB1 rescue studies, CRISPR-mediated knockout of *Lkb1* in murine KP NSCLC cells led to increased cell surface expression of CD38 (**Fig. 3B, 3C, Supplementary Fig. S4A**). LKB1 loss also induced *Sik1* mRNA expression, consistent with the aforementioned feedback circuit for *Sik1* transcriptional regulation (30) (**Supplementary Fig. S4B-S4D, Supplementary Table S5**). We subsequently tested the impact of knockout of *Sik1*, *Sik2* and *Sik3*, alone or in combination. We observed prominent increases in CD38 expression at the cell surface in KP cells upon inactivation of *Sik1* or of all three SIK family members (**Fig. 3B, 3C, Supplementary Fig. S4B-S4D**). LKB1 or SIK loss also upregulated total CD38 protein levels and induced CD38 mRNA expression as determined by western blot and real-time PCR analysis, respectively (**Fig. 3D, 3E**). Likewise, acute pharmacologic inhibition of SIK kinase activity with the pan-SIK inhibitor, YKL-05-099 (31), upregulated CD38 in multiple LKB1 WT mouse cancer cell lines (**Supplementary Fig. S4E**). These data indicate that LKB1 suppresses CD38 expression level through SIK family kinases, mainly through SIK1. Since the increase in CD38 levels upon SIK inactivation did not reach that seen in KP sg*Lkb1* cells (**Fig. 3B, 3C)**, additional factors downstream of LKB1 might also contribute to the suppression of CD38 expression. Notably, the tumorigenicity of KP cell lines was enhanced by either *Lkb1* or *Sik1/2/3* knockout, as demonstrated by orthotopic allograft models, producing a lung tumor burden comparable to tumors from KL mouse lung cancer cell lines (**Fig. 3F, 3G**). Moreover, as in the case of allografts with LKB1 inactivation, *Sik1/2/3*-KO NSCLC allografts showed significant upregulation of ADP ribose, increase of which was the single outlier among 147 total metabolites tested (**Fig. 3H**)

The CREB Regulated Transcription Coactivator family (CRTC1, CRTC2, and CRTC3) proteins are known targets of SIK. SIK-mediated phosphorylation of CRTCs at a 14-3-3 motif leads to their cytoplasmic retention. Loss of this phosphorylation due to LKB1 or SIK inactivation causes CRTC nuclear translocation and complex formation with cAMP response element binding protein (CREB) and CREB binding protein (CBP/p300) to initiate gene transcription through cAMP response element (CRE) binding sites (32–35). Analysis of the CREB transcriptional activity using a reporter with a synthetic CRE motif (36) demonstrated that LKB1 or SIK1/2/3 loss in murine KP tumor cells resulted in markedly increased CREB reporter activity (**Fig. 3I**). Moreover, analysis of the ENCODE ChIP-seq database (37) revealed binding of p300 (a.k.a. EP300) to the CD38 promoter at the CRE site (**Fig. 3J**). These results are consistent with CRTC/CREB/P300 activation mediating CD38 induction upon LKB1 or SIK loss of function (**Supplementary Fig. S5A**). Collectively, the data indicate that inactivation of LKB1/SIK signaling drives CREB-mediated transcriptional upregulation of the NAD-consuming enzyme CD38 *in vivo*.

### LKB1 mutant tumor models are sensitive to anti-CD38 antibodies

We next sought to test whether the increase in CD38 in LKB1 mutant cancers confers an emergent vulnerability that could be exploited therapeutically. Multiple drugs targeting CD38, including the monoclonal antibodies daratumumab and isatuximab, are approved for use in multiple myeloma (MM), a plasma cell malignancy characterized by high cell surface CD38 expression (21,22). Daratumumab is the first approved antibody against CD38, which is a fully human IgG1-κ monoclonal antibody that kills MM cells through multiple mechanisms, including antibody-dependent cellular cytotoxicity (ADCC) via binding to Fcγ receptors (FcγR) (38–40). We first performed *in vitro* cytotoxicity assays by co-culturing the natural killer cell line NK-92 with a series of *LKB1* mutant (A549, HCC44) and *LKB1* WT (H358, H441) NSCLC cell lines in the presence or absence of daratumumab. In A549 cells, daratumumab-mediated cytotoxicity was observed only in the context of co-culture with NK-92 (**Fig. 4A**), whereas HCC44 showed a response to daratumumab alone, which was enhanced by NK cell co-culture (**Fig. 4B**). Conversely, *LKB1* WT cell lines (H358 and H441) did not exhibit significant cytotoxic responses to daratumumab treatment, irrespective of NK cell presence (**Fig. 4C, Supplementary Fig. S5B**). There also were dose-dependent increases in daratumumab-induced ADCC in the *LKB1* mutant cell lines but not in the *LKB1* wild type H358 cell line (**Fig. 4D**).

**Figure 4.**
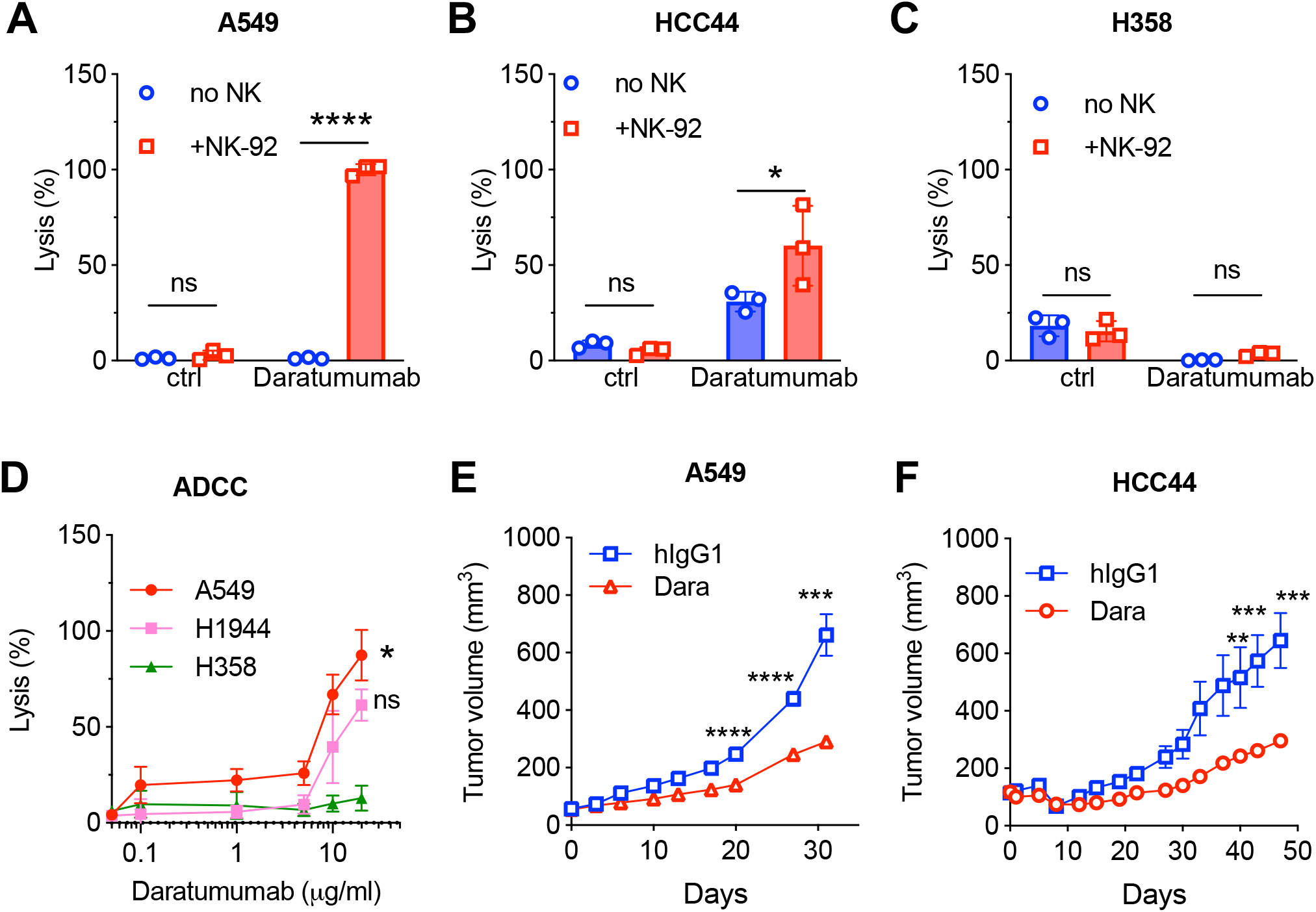
Daratumumab induces killing of CD38 positive lung cancer cells through antibody-dependent cellular cytotoxicity (ADCC). **A-B,** Cytotoxicity assay in LKB1 mutant A549 (**A**), HCC44 (**B**) or LKB1 wild type H358 (**C**) target cells using calcein-acetoxymethyl (AM) release in the presence of NK cell line NK-92. Lytic activity was measured 4 hours after co-culturing (E:T=100:1) in the presence of Daratumumab (20 μg/ml). Student’s *t* test, two-tailed analysis was performed to compare with no NK cell control as indicated. **D**, Indicated cells were cultured with NK-92 cells (E:T=100:1) in the presence of increasing concentrations of daratumumab. The percentage of cell lysis for each cell line was measured 4 hrs after co-culturing. Paired student’s *t* test for LKB1 mutant cells (A549 or H1944) compared with LKB1 wild type control H358. **E-F,** Mouse A549 (**E**) or HCC44 (**F**) xenograft tumor growth in response to daratumumab treatment. For each mouse, 3x10^6^ A549 cells (**E**) or 5x10^5^ HCC44 (**F**) were implanted into nu/nu mice subcutaneously. Daratumumab or hIgG1 (12.5mg/kg, once per week) was dosed when tumor size reached100 mm^3^. Multiple unpaired student’s *t* test was performed for each of the time points indicated. \**P*<0.05, \*\**P*<0.01, \*\*\**P*<0.001, \*\*\*\**P*<0.0001.

To study the effects of daratumumab *in vivo*, we implanted the LKB1 mutant A549 and HCC44 cell lines into athymic nude (nu/nu) mice to generate tumor xenografts. Daratumumab is a human IgG1 that is recognized by mouse FcγRs on effector cells, resulting in ADCC via crosstalk with mouse NK cells (41). This enables evaluation of NK cell-mediated tumor cell killing *in vivo*. When tumors reached approximately 100 mm^3^ in volume, mice were treated with daratumumab once weekly. We observed a significant reduction in tumor growth in both mutant LKB1 tumor models (**Fig. 4E, 4F, Supplementary Fig. S5C-S5F**), supporting the potential of harnessing CD38 as a target in these tumors.

## DISCUSSION

LKB1 as one of the most commonly mutated genes in NSCLC. Despite extensive studies of the biology of *STK11*-mutant NSCLC, there are presently no effective targeted therapies for this aggressive subtype. LKB1 reprograms cellular metabolism through AMPK-related kinases and plays a pleiotropic role in tumor growth, metastasis, and immune evasion. Here, we sought to uncover metabolic vulnerabilities of *STK11*-mutant NSCLC by in vivo metabolomic analysis. These studies led us to identify CD38 as being highly overexpressed in these tumors. Importantly, preclinical studies using an the approved anti-CD38 antibody, daratumumab, suggested the potentially of exploiting CD38 overexpression therapeutically.

Our work reveals new insights into the metabolism of LKB1 mutant tumors *in vivo*. By performing unbiased metabolomics analysis of KL versus KP tumors, we observed significant increases of ADP-ribose and glutamate upon LKB1 loss. ADP-ribose is generated by multiple NAD^+^ consuming enzymes, including PARPs, sirtuins and ectoenzyme CD38. Among the NAD+ consuming enzymes, we discovered that only CD38 was upregulated enzyme upon LKB1 loss-of-function. The CD38 ectoenzyme plays a central role in tissue NAD^+^ homeostasis, serving as a major NAD glycohydrolase and ADP-ribosyl cyclase that generate second messengers ADP-ribose and cyclic ADP-ribose (cADPR). Notably, we discovered that SIK family kinases are responsible for LKB1-mediated suppression of CD38 expression. This is through the transcriptional regulation of CREB binding site at the promoter of CD38. LKB1/SIK loss-of-function leads to dephosphorylation and nuclear translocation of CRTCs, and subsequently, the formation of CRTC/CREB/CBP/p300 complex that initiates transcription via CRE binding sites. Consistently, the ChIP-seq data confirmed that p300 binds to the CRE site of CD38 promoter on LKB1 mutant NSCLC cancer cells. As a result, the expression of CD38 on the cell surface increased ∼5-50 fold in various lung cancer cells with LKB1 mutation.

CD38 is highly expressed in multiple myeloma, and several therapeutic antibodies and ADCs against CD38 are FDA approved for the treatment of this disease, including daratumumab and isatuximab (42,43). In addition, CD38 is also implicated in immune suppression. ADP-ribose generated by CD38-mediated hydrolysis of NAD+ to ADP-ribose is further broken down by CD203a and CD73, yielding adenosine. The binding of adenosine binding to its receptors (A2AR or A2BR) on different immune lineages leads to diverse immune suppressive processes (44–46). The CD38-CD203a-CD73 pathway can contribute to breast cancer relapse, whereas combined inhibition of NAD generation and of the adenosine receptor reactivates CD8+ T cells (47,48). Notably, LKB1 mutant cancers are highly immune suppressive and resistant to immune checkpoint blockade (9,23,49). Effects of CD38 on both the NAD+ salvage and adenosine pathways could drive metabolic adaptation and immune suppression in these cancers.

As a clinically approved drug, anti-CD38 antibody daratumumab did not show dose-limiting toxic events up to 24 mg/kg and no maximum tolerated dose was found in a phase 2 clinical trial on multiple myeloma patients (21). Thus, targeting CD38 in LKB1 mutant cancer patients may likewise have minimal side effects or toxicity. Instead, inhibiting CD38 in the tumor microenvironment will potentially induce anti-tumor immunity through the suppression of adenosine pathway via the CD38-CD73 axis. Due to limitation in the availability of CD38 neutralizing antibody against the murine protein, we are not able to further validate the effect of impact of targeting CD38 on the immune system of LKB1 mutant lung cancers.

Our study identified a mechanistic link between loss of function in LKB1-SIK signaling and transcriptional upregulation of CD38. The resulting increase of CD38 expression at the cell correlated with an elevation in the product of NAD^+^ metabolism, ADP-ribose. Importantly, *in vivo* treatment of LKB1 mutant lung tumors with a CD38 targeting monoclonal antibody arrested tumor growth. Thus, CD38 is an emergent vulnerability unique to LKB1 mutant NSCLC and is a promising therapeutic target.

## METHODS

### Metabolomic Profiling

Mouse lung tumor nodules were resected, snap frozen in liquid nitrogen, and processed at NYU Langone Health Metabolomics Core for hybrid metabolomics profiling. Prior to extraction, samples were moved from -80 °C storage to wet ice and thawed. Extraction buffer, consisting of 80% methanol (Fisher Scientific) and 500 nM metabolomics amino acid mix standard (Cambridge Isotope Laboratories, Inc.), was prepared and placed on dry ice. Samples were extracted by mixing with a ratio of 10 mg of sample with 1000 µL of extraction buffer in 2.0mL screw cap vials containing ∼100 µL of disruption beads (Research Products International, Mount Prospect, IL). Each was homogenized for 10 cycles on a bead blaster homogenizer (Benchmark Scientific, Edison, NJ). Cycling consisted of a 30 sec homogenization time at 6 m/s followed by a 30 sec pause. Samples were subsequently spun at 21,000 g for 3 min at 4 °C. A set volume of each (270 µL) was transferred to a 1.5 mL tube and dried down by speedvac (Thermo Fisher, Waltham, MA). Samples were reconstituted in 30 µL of Optima LC/MS grade water (Fisher Scientific, Waltham, MA). Samples were sonicated for 2 mins, then spun at 21,000g for 3 mins at 4°C. 20 µL were transferred to LC vials containing glass inserts for analysis. The remaining sample was placed in -80 °C for long term storage.

### Metabolomics Data Processing

The resulting Thermo^TM^ RAW files were converted to SQLite format using an in-house python script to enable peak detection and quantification. The centroided data were searched using an in-house python script and peak heights were extracted from the SQLite file based on a previously established library of metabolite retention times and accurate masses adapted from the Whitehead Institute (50), and verified with authentic standards and/or high resolution MS/MS spectral manually curated against the NIST14MS/MS and METLIN (2017) tandem mass spectral libraries (51,52). Metabolite peaks were extracted based on the theoretical *m*/*z* of the expected ion type e.g., [M+H]^+^, with a ±15 part-per-million (ppm) tolerance, and a ± 7.5 second peak apex retention time tolerance within an initial retention time search window of ± 0.5 min across the study samples. The resulting data matrix of metabolite intensities for all samples and blank controls was processed with an in-house statistical pipeline Metabolyze version 1.0 and final peak detection was calculated based on a signal to noise ratio (S/N) of 3X compared to blank controls, with a floor of 10,000 (arbitrary units). For samples where the peak intensity was lower than the blank threshold, metabolites were annotated as not detected, and the threshold value was imputed for any statistical comparisons to enable an estimate of the fold change as applicable. The resulting blank corrected data matrix was then used for all group-wise comparisons, and t-tests were performed with the Python SciPy (1.1.0) library to test for differences and generate statistics for downstream analyses. Any metabolite with p-value < 0.05 was considered significantly regulated (up or down). The final heatmap with selected significant metabolites was generated by performing row-wise unsupervised hierarchical clustering and column-wise supervised clustering on the imputed matrix values utilizing the R library pheatmap (1.0.12).

### Cell Culture

Mouse lung cancer cell lines were cultured in RPMI 1640 (Gibco, Thermo Fisher Scientific) supplemented with 10% fetal bovine serum (FBS, Gibco) and antibiotics. The *Kras*^LSL-G12D/+^*Trp53*^fl/fl*-/-*^ (KP) and *Kras*^LSL-G12D/+^*Lkb1*^fl/fl^ (KL) murine lung cancer cell lines were generated and benchmarked as previously described^4^. Briefly, lung nodules from KP and KL GEMMs were dissociated to single cell suspensions with collagenase D and DNase I digestion, seeded on tissue culture plates, and cultured in RPMI 1640 medium supplemented with 10% FBS and antibiotics. FC4662 mouse pancreatic cancer and were cultured in DMEM supplemented with 10% FBS and antibiotics. Human NSCLC cell lines H1944, HCC44, A549, A427, H441, H358, Calu1 and H1792 were cultured with RPMI1640 supplemented with 10% FBS and antibiotics. HEK-293 cells were obtained from ATCC and cultured in DMEM (Gibco) supplemented with 10% FBS. Human isogenic lines A549 pBABE, A549-LKB1, A549-LKB1-KD were cultured in RPMI 1640 supplemented with 10% FBS and puromycin (5 μg/ml). All cells were seeded in T75 flask and cultured at 37°C in a humidified incubator at 5% CO_2_ and verified on a bi-weekly basis to be mycoplasma negative.

### Plasmid construction and lentiviral generation and transduction

Plasmids pLenti-Cas9-puro and psPAX2 and pMD2.G were purchased from Addgene. The sgRNAs specific for mouse *Sik1*, *Sik2*, *Sik3* and *Lkb1* were cloned into pXPR-RFP-Blast using the Gibson Assembly Kit (E2611L, NEB). The target sequences are as follows: *Sik1* sgRNA: 5’-CTCCTACCCAGACGTCCAGC-3’, *Sik2* sgRNA: 5’-ATCCTAATAGATTTCGGCTT-3’, *Sik3* sgRNA: 5’-GCCGCCCCAGAGCTCTTCGA-3’, *Lkb1*sgRNA: 5’-CGAGACCTTATGCCGCAGGG-3’. vector using primer sequences listed the methods from our prior publication^5^. The CREB reporter pLminP_Luc2P_RE16 was purchased from Addgene (Cat # 90357).

Stable cell lines were generated using lentiviral transduction, which were first generated by co-transfecting packaging vector psPAX2, envelope vector pMD2.G, and the expression vectors into HEK-293 cells using Lipofectamine 3000. Transfection medium was removed and HEK-293 cells were cultured in RPMI 1640 + 10% FBS for 48 hours. Viruses were then syringe-filtered through a 0.45 µm filter (corning) to remove debris. The filtered viruses were added to culture media of cells with polybrene (Santa Cruz, final concentration of 8 µg/mL). Fresh media were replaced 48 hours after infection, and infected cells expressing the constructs were selected with puromycin and blasticidin with optimized concentration for each cell line with different gRNA. For CREB reporter expression stable cell lines, cells were sorted with GFP+ and RFP+ 48 hrs after infection with CREB expression lentivirus and cultured in RPMI media for future analysis.

### Western blotting and antibodies

Cells were washed twice with ice-cold PBS, scraped and collected as pellets post centrifugation at 3000x r.p.m. for 5 min. The collected cell pellets were lysed with RIPA lysis buffer (ThermoFisher Scientific cat# 89900) supplemented with protease and phosphatase inhibitor cocktail (ThermoFisher Scientific Cat# 78440). Protein concentrations were quantified using BCA protein assay kit (ThermoFisher Scientific cat# 23225) with bovine serum albumin (BSA) as the protein standard. Equal amount of protein lysates were boiled in 5x SDS sample buffer with bromophenol blue (Boston BioProducts cat#BP-111R) for 7 minutes and separated with 4-20% Mini-PROTEAN TGX protein Gel (Bio-Rad, Cat # 5671094), transferred to nitrocellulose membranes according to standard protocol. Membranes were blocked at room temperature with TBS blocking buffer (LI-COR, Cat # 927-60001), and incubated with corresponding primary antibodies overnight at 4 °C with standard protocol. Secondary antibody of IRDye 680RD donkey anti-Mouse IgG (Li-COR cat# 925-68072) and IRDye 800CW donkey anti-rabbit IgG (LI-COR cat# 925-32212) were used and the fluorescent signal on the membrane were imaged on Odyssey classic infrared imaging system (LI-COR) using Image Studio Lite (V 5.2). Primary antibodies used are human CD38 (Cell Signaling Technology, Cat #51000S), mouse CD38 (Cell Signaling Technology, Cat #92457), LKB1 (Cell Signaling Technology, Cat #3047), β-actin (Sigma, Cat# A5441).

### RNA extraction and quantitative RT-PCR

Total RNA was extracted using RNeasy Mini Kit (Qiagen) according to manufacturer protocol, and cDNA was generated using High-Capacity RNA-to-cDNA kit (ThermoFisher Scientific, Cat# 4387406). Quantitative PCR with reverse transcription (qRT-PCR) assays were performed using PowerUp SYBR Green Master Mix (ThermoFisher Scientific Cat# A25742) on the StepOne Real-Time PCR System (ThermoFisher Scientific) following the manufacturer’s instructions. Results were analyzed using the ΔΔ*C*_t_ method by normalizing with *Actb* reference gene. Primer sequences are listed in Extended Data Table 5.

### Flow Cytometry

Tumors were mechanically dissociated followed by enzymatic digestion in HBSS containing 2 U/mL DNAse and 0.5% w/v collagenase type D at 37°C for 45 minutes and tumor cell suspensions were filtered through a 70 μm cell strainer (Fisher Scientific). All cell suspensions were centrifuged at 1500x rpm for 5 minutes and resuspended in RBC lysis buffer (Biolegend) for 5 minutes at room temperature. RBC lysis was halted by the addition of PBS and cells were centrifuged and resuspended in FACS buffer (PBS + 2% FBS). Tumor cell suspensions were counted and 2 x 10^6^ tumor cells per sample were probed with the indicated fluorescently labeled antibodies. Cells were then washed and stained with indicated fluorescently labeled antibodies. Cell lines were trypsinized after culturing, filtered through 70 μm cell strainer, and washed with 1x PBS. After centrifugation, the cell pellets were stained with live/dead fixable dye (ThermoFisher Scientific, Cat # L34966) followed by staining with indicated fluorescently labeled antibodies with standard protocol and FACS analysis were performed on BD Fortessa LSR flow cytometer. FlowJo 10.6.1 was used to analyze FACS data. Flow antibodies used are anti-mouse CD38 (clone 90, Biolegend), anti-mouse EpCAM (clone G8.8), anti-mouse CD45 (clone 30-F11), anti-mouse CD31 (clone 390), anti-human CD38 (clone HIT2), anti-human EpCAM (clone 9C4).

### Patients IHC

Human lung tumor specimens were collected in Fudan University Shanghai Cancer Center, with patient written consents and the approval from the Institute Research Ethics Committee. All tumor specimens were taken at the time of surgical resection. And all cases were re-reviewed by pathologists for confirmation of tumor histology. In the present study, 17 human lung ADC samples were performed IHC staining with an anti-CD38 antibody (Cell Signaling Technology, Cat #51000).

### Lung Cancer Patient Transcriptomic Analysis

Human lung adenocarcinoma RNA-seq data were obtained from The Cancer Genome Atlas (TCGA) publicly available database. Patient datasets with *KRAS* mutations were further stratified into groups with *TP53* (KP) or *STK11/LKB1* (KL) co-occurring mutations for downstream comparative analyses. For specific gene expression analysis, TCGA RNAseq files from KL and KP human NSCLC were obtained through the GDC portal. TPM values for analyzed genes from each patient sample were compared between KL and KP cohorts.

### CREB reporter assay

The The CREB reporter pLminP_Luc2P_RE16 was purchased from Addgene and lentivirus were produced as described above. After virus transduction into target cells of KP gRNA lines as indicated, infected cells were sorted as GFP+ cells and cultured in culture media. The promoter activity for each cell line was determined using Steadylite Plus reporter Gene Assay System (PerkinElmer Cat # 6066751) according to manufactural manual. Briefly, cells were plated onto 96-well plate overnight, and changed to HBSS with calcium and magnesium (ThermoFisher Scientific, Cat # 14025092) (100 μl/well). Then 100 μl of reconstituted steadylite plus buffer were added to each well and the luminescence were measured using Synergy Neo2 Hybrid Multi-Mode Microplate Reader (Agilent) within 4 hrs.

### ChIP-seq data analysis

The ChIP-seq data of A549 with EP300 were obtained from Encyclopedia of DNA Elements (ENCODE) Project dataset with transcriptional factor ChIP-seq (TF ChIP) pipeline (accession # ENCSR886OEO and ENCSR686BQM) ^6^. The sequenced data were analyzed with ENCODE4 (v1.1.6) and aligned with GRCh38 genome assembly and annotation.

### Antibody directed cell cytotoxicity (ADCC)

Lysis of tumor cells by ADCC was measured using Calcein-AM release assay as previously described ^7,8^. Briefly, cells were washed once with 1x PBS and incubated with 10 μM calcein-AM (Biolegend Cat # 425201) for 30 min at 37°C in the incubator. After incubation, cells were washed twice with culture medium and plated overnight. Then NK-92 cells were added to the indicated tumor cells with effector:target (E:T) ratior of 100:1 in the presence of different concentrations of daratumumab as indicated. The percentage of specific lysis was determined according to the formula [(test release-spontaneous release)/maximum release – spontaneous release]]X100 as previously described^7,8^. Spontaneous release represents Calcein-AM release in media alone, and maximum release were determined by adding 2% Triton X-100 to medium for target cells.

### Mice Experiments

All animal experiments were reviewed and approved by the Institutional Animal Care and Use Committee (IACUC) at NYU Langone Health. For lung tumor growth assays, 10^6^ KP or KP-gRNA syngeneic mouse lung cancer cells were implanted into male C57BL/6 mice at 7 weeks of age via tail vein injection. KL cells were injected into female C57BL/6 mice at 1x10^6^/mouse via tail vein injection. After 6 weeks, mice were imaged by MRI to assess tumor volume. For xenograft model, A549 (3x10^6^/tumor/mouse) or HCC44 (5x10^5^/tumor/mouse) were mixed with Matrigel (1:1 ratio, 200ul each tumor) and implanted into nu/nu mice subcutaneously.

Mouse weights were measured weekly to adjust total dosage and assess the effects of drug combinations on mouse health. After euthanasia by CO_2_ exposure at 3 L/min, syngeneic primary tumors and/or mouse lungs were formalin-fixed, paraffin-embedded, and sectioned for histological analysis.

## Data availability

The data supporting the findings of this study are available within the paper. Raw data of metabolomics analysis for this study were generated at NYU Metabolomics Core Resource Laboratory. The RNAseq data data analyzed in this study were obtained from Gene Expression Omnibus (GEO) at GSE137244 and GSE137396. TCGA data used is publicly available at GDC portal (https://portal.gdc.cancer.gov/) and patient sample numbers used are provided in the paper (Table S1). ChIPseq data is available from Encyclopedia of DNA Elements (ENCODE) Project dataset (https://www.encodeproject.org). The data were derived from A549 ChIPseq (TF ChIP) results with accession number ENCSR886OEO and ENCSR686BQM. Proteomics and transcripts raw datasets of lung cancer patients with LKB1 mutation are publicly available through the CPTAC data portal (https://pdc.cancer.gov/pdc/) study ID PDC000149, and PDC portal (https://portal.gdc.cancer.gov) with dbGaP Study Accession: phs001287.v5.p4, respectively. All other data supporting the findings of this study are available from the corresponding author upon request.

## Notes

**Financial support:** This work is supported by R01CA248896 (B.G.N. and K.K.W.) and P30CA016087 (Cancer Center Support Grant, B.G.N.).

**Disclosure statement of conflict of interest**: K.K.W. is a founder and equity holder of G1 Therapeutics and has sponsored research agreements with Takeda, TargImmune, Bristol-Myers Squibb (BMS), Mirati, Merus, Alkermes, and consulting and sponsored research agreements with AstraZeneca, Janssen, Pfizer, Novartis, Merck, Zentalis, BridgeBio and Blueprint. C.M.R. reports personal fees from AbbVie, Amgen, Ascentage, AstraZeneca, Bicycle, Celgene, Daiichi Sankyo, Genentech/Roche, Ipsen, Jansen, Jazz, Lilly/Loxo, Pfizer, PharmaMar, Syros, Vavotek, Bridge Medicines and Harpoon Therapeutics, outside the submitted work. B.G.N. reports other support from NYU Langone Health, Northern Biologics, and Navire Pharma, personal fees and other support from Lighthorse Therapeutics, Arvinas, Recursion Pharma, GLG Group, and Repare Therapeutics, and grants from the NCI/NIH (R21CA267362, R01CA248896, and P30CA016087) during the conduct of the study; a patent for 63402606 pending; and during the course of performing these experiments, his wife owned personal shares in Amgen and Regeneron. B.G.N. also own shares of Mirati. B.G.N. is the co-founder, hold equity and receive consulting revenue from Aethon Therapeutics. A.N.H. has received research support from Amgen, Blueprint Medicines, BridgeBio, Bristol-Myers Squibb, C4 Therapeutics, Eli Lilly, Nuvalent, Pfizer, Roche/Genentech, Scorpion Therapeutics; has served as a compensated consultant for Nuvalent, Tolremo Therapeutics, Engine Biosciences and TigaTx. S.K. reports grants from Puretech Health and Argenx BVBA, and grants and personal fees from Black Diamond Therapeutics outside the submitted work; a patent for PCT/US2022/018171 pending; and is a cofounder of and holds equity in Revalia Bio.

### Competing Interest Statement

K.K.W. is a founder and equity holder of G1 Therapeutics and has sponsored research agreements with Takeda, TargImmune, Bristol-Myers Squibb (BMS), Mirati, Merus, Alkermes, and consulting and sponsored research agreements with AstraZeneca, Janssen, Pfizer, Novartis, Merck, Zentalis, BridgeBio and Blueprint. C.M.R. reports personal fees from AbbVie, Amgen, Ascentage, AstraZeneca, Bicycle, Celgene, Daiichi Sankyo, Genentech/Roche, Ipsen, Jansen, Jazz, Lilly/Loxo, Pfizer, PharmaMar, Syros, Vavotek, Bridge Medicines and Harpoon Therapeutics, outside the submitted work. B.G.N. reports other support from NYU Langone Health, Northern Biologics, and Navire Pharma, personal fees and other support from Lighthorse Therapeutics, Arvinas, Recursion Pharma, GLG Group, and Repare Therapeutics, and grants from the NCI/NIH (R21CA267362, R01CA248896, and P30CA016087) during the conduct of the study; a patent for 63402606 pending; and during the course of performing these experiments, his wife owned personal shares in Amgen and Regeneron. B.G.N. also own shares of Mirati. B.G.N. is the co-founder, hold equity and receive consulting revenue from Aethon Therapeutics. A.N.H. has received research support from Amgen, Blueprint Medicines, BridgeBio, Bristol-Myers Squibb, C4 Therapeutics, Eli Lilly, Nuvalent, Pfizer, Roche/Genentech, Scorpion Therapeutics; has served as a compensated consultant for Nuvalent, Tolremo Therapeutics, Engine Biosciences and TigaTx. S.K. reports grants from Puretech Health and Argenx BVBA, and grants and personal fees from Black Diamond Therapeutics outside the submitted work; a patent for PCT/US2022/018171 pending; and is a cofounder of and holds equity in Revalia Bio.

