## Supplementary material for "*In vivo* metabolomics identifies CD38 as an emergent vulnerability in *LKB1*-mutant lung cancer": Table 1

**Table 1. Metabolite measurement from KL and KP tumors**

| Metabolite | Formula | KP #1 | KP #2 | KP #3 | KP sgSIK1/2/3 #1 | KP sgSIK1/2/3 #2 | KP sgSIK1/2/3 #2 | KL #1 | KL #2 |
|---|---|---|---|---|---|---|---|---|---|
| L-Phenylalanine | C9H11NO2 | 108083200 | 115886080 | 121387744 | 136953744 | 96578480 | 101577216 | 65568744 | 70944496 |
| Uracil | C4H4N2O2 | 2870513 | 3167644 | 2494798 | 1824128 | 1357097 | 4273245 | 1017940 | 680633 |
| Adenosine diphosphate ribose | C15H23N5O14P2 | 813273 | 897648 | 921325 | 1111101 | 1467367 | 1596786 | 1687757 | 2240976 |
| L-Tryptophan | C11H12N2O2 | 8132383 | 12290281 | 11881540 | 7955288 | 12982164 | 4085455 | 2520534 | 4682062 |
| Pipecolic acid | C6H11NO2 | 67083852 | 46075500 | 59731936 | 130931632 | 18452330 | 42079084 | 19083070 | 31620676 |
| Citrulline | C6H13N3O3 | 198119200 | 187677568 | 177471744 | 148029232 | 96311384 | 129865400 | 58343896 | 43733608 |
| L-Proline | C5H9NO2 | 513824608 | 536034560 | 618536896 | 626748096 | 440863072 | 466742432 | 264769808 | 336892352 |
| L-Valine | C5H11NO2 | 190701360 | 195815568 | 129985984 | 194125184 | 94665560 | 124038088 | 96124600 | 99246840 |
| Acetoacetic acid | C4H6O3 | 471498 | 533155 | 731421 | 342808 | 1172516 | 907292 | 1623317 | 954473 |
| L-Glutamic acid | C5H9NO4 | 254452864 | 277129440 | 273437696 | 221834304 | 285253888 | 220148912 | 285660608 | 297457152 |
| Allantoin | C4H6N4O3 | 3859260 | 3560545 | 2282092 | 3352287 | 477195 | 2286959 | 733653 | 1505631 |
| L-Methionine | C5H11NO2S | 108946872 | 106572464 | 182398176 | 178340816 | 96646056 | 148268368 | 50171180 | 50347412 |
| Ureidopropionic acid | C4H8N2O3 | 3830996 | 3462592 | 4910071 | 4341748 | 4730302 | 4158682 | 2089698 | 1974821 |
| L-Threonine | C4H9NO3 | 9007562 | 8608225 | 10421532 | 16050596 | 7216238 | 9307981 | 4014280 | 4858095 |
| L-Carnitine | C7H16NO3 | 455245312 | 482876224 | 649808896 | 359054688 | 807283584 | 673899392 | 1032814976 | 678722816 |
| L-Isoleucine | C6H13NO2 | 174216512 | 177747584 | 112021408 | 126848408 | 117756704 | 131244808 | 117723864 | 75696792 |
| Cyclic AMP | C10H12N5O6P | 32313 | 32158 | 33557 | 15577 | 27233 | 18732 | 24779 | 21308 |
| L-Leucine | C6H13NO2 | 167020720 | 131213680 | 109830168 | 120031424 | 116331800 | 130143016 | 110828616 | 68096704 |
| L-Tyrosine | C9H11NO3 | 23137578 | 27755160 | 26751714 | 19890380 | 31801028 | 23289930 | 18187538 | 13955781 |
| Uridine | C9H12N2O6 | 5224356 | 3546023 | 2477575 | 2454234 | 1683358 | 2888180 | 2581953 | 1798642 |
| N-Acetylserine | C5H9NO4 | 9365471 | 6104942 | 7204762 | 4382895 | 5280302 | 3496178 | 4353554 | 6385790 |
| NAD | C21H28N7O14P2 | 111236 | 373245 | 200201 | 0 | 246944 | 0 | 14007 | 120467 |
| Butyrylcarnitine | C11H21NO4 | 12286415 | 14854061 | 20453914 | 15374136 | 28673084 | 19122408 | 25218328 | 17494010 |
| Hypoxanthine | C5H4N4O | 275903040 | 324877920 | 224132816 | 142407024 | 161504944 | 247218688 | 236754656 | 130949744 |
| N-Acetylornithine | C7H14N2O3 | 6693790 | 5732840 | 7998640 | 8003153 | 10193081 | 7750960 | 5952694 | 3501154 |
| N-Acetylglutamine | C7H12N2O4 | 189831 | 126258 | 78885 | 103016 | 79435 | 101418 | 74507 | 64260 |
| Phosphorylcholine | C5H14NO4P | 959584000 | 1084129920 | 562594624 | 720667712 | 516838912 | 605986624 | 473807584 | 599282048 |
| Pyruvic acid | C3H4O3 | 341765 | 382625 | 191731 | 181490 | 377093 | 233010 | 387170 | 724374 |
| Argininosuccinic acid | C10H18N4O6 | 36817 | 30453 | 23691 | 24064 | 24389 | 28917 | 16723 | 25614 |
| Carnosine | C9H14N4O3 | 201930 | 147684 | 126471 | 155759 | 105998 | 143952 | 174122 | 281756 |
| S-Adenosylhomocysteine | C14H20N6O5S | 265879 | 501888 | 242667 | 316493 | 127524 | 303527 | 171408 | 244549 |
| N-Acetyl-L-phenylalanine | C11H13NO3 | 1270170 | 875428 | 437446 | 823302 | 401233 | 513353 | 191497 | 408080 |
| L-Alanine | C3H7NO2 | 81104016 | 94439312 | 114504224 | 99014104 | 109581664 | 66682244 | 47324244 | 68837104 |
| Betaine | C5H11NO2 | 947436800 | 890990464 | 1115118976 | 609426752 | 1094455040 | 862793472 | 1174147840 | 1041329472 |
| Phosphoserine | C3H8NO6P | 30644 | 40002 | 36327 | 31897 | 28528 | 24235 | 19394 | 33886 |
| 3-Phosphoglyceric acid | C3H7O7P | 419153 | 757571 | 79447 | 67900 | 217135 | 76567 | 100401 | 86299 |
| Guanosine | C10H13N5O5 | 1602226 | 959686 | 325267 | 257158 | 333624 | 296361 | 464087 | 252605 |
| Fructose 6-phosphate | C6H13O9P | 393044 | 472005 | 254333 | 449160 | 485523 | 1532731 | 3092662 | 924334 |
| Aminoadipic acid | C6H11NO4 | 789850 | 589284 | 544526 | 515185 | 324301 | 396030 | 454737 | 617127 |
| Phosphocreatine | C4H10N3O5P | 2596263 | 565055 | 404004 | 1251753 | 272260 | 53800 | 59816 | 25299 |
| Inosinic acid | C10H13N4O8P | 2523573 | 2548241 | 2840408 | 673842 | 848476 | 1451590 | 3406744 | 3433935 |
| Guanidoacetic acid | C3H7N3O2 | 26664966 | 20556382 | 67723768 | 20167816 | 23577784 | 24293816 | 19010614 | 15363747 |
| Glucose 6-phosphate | C6H13O9P | 612504 | 603968 | 706192 | 569009 | 777991 | 1751217 | 3438194 | 1106820 |
| L-Lactic acid | C3H6O3 | 141622416 | 166370416 | 150449776 | 142671696 | 293589536 | 241324496 | 124988240 | 134175088 |
| L-Histidine | C6H9N3O2 | 18994582 | 7936852 | 16739186 | 41154104 | 22353980 | 17575388 | 9918621 | 5927307 |
| L-Acetylcarnitine | C9H17NO4 | 420369408 | 458540992 | 726088256 | 275137184 | 788141120 | 496303776 | 841848256 | 550785152 |
| O-Phosphoethanolamine | C2H8NO4P | 27114428 | 39253484 | 24351066 | 23017610 | 26081174 | 17042724 | 22048898 | 24635310 |
| Xanthine | C5H4N4O2 | 12287432 | 13990865 | 7523732 | 6208266 | 5964422 | 15871897 | 10063852 | 5208152 |
| Uridine diphosphate glucose | C15H24N2O17P2 | 1298136 | 1547095 | 1079635 | 1271972 | 874449 | 871755 | 1586988 | 2137216 |
| Phosphoenolpyruvic acid | C3H5O6P | 283999 | 429975 | 43478 | 31770 | 121889 | 81406 | 23497 | 79649 |
| Xanthosine | C10H12N4O6 | 33020 | 40612 | 14224 | 11801 | 3810 | 56585 | 26497 | 4866 |
| cis-Aconitic acid | C6H6O6 | 4660194 | 3840221 | 2673810 | 2462363 | 2114080 | 2391391 | 2550276 | 2806956 |
| ADP | C10H15N5O10P2 | 1698092 | 4377490 | 892778 | 343030 | 1061271 | 393034 | 564926 | 1374806 |
| 5'-Methylthioadenosine | C11H15N5O3S | 8876233 | 10706533 | 12049495 | 11350584 | 10332823 | 12365457 | 8276610 | 9091155 |
| L-Asparagine | C4H8N2O3 | 16470313 | 15280018 | 26281424 | 20035002 | 25899462 | 20027726 | 12961705 | 12281071 |
| 6-Phosphogluconic acid | C6H13O10P | 42606 | 40542 | 16028 | 158683 | 39831 | 25603 | 86323 | 83433 |
| L-Kynurenine | C10H12N2O3 | 648335 | 621682 | 1891286 | 1617436 | 859190 | 11379712 | 2573112 | 751172 |
| Ribose 1-phosphate | C5H11O8P | 1955607 | 922470 | 957209 | 1159915 | 1433369 | 1301482 | 2459848 | 1881183 |
| CDP | C9H15N3O11P2 | 24571 | 45657 | 10729 | 6880 | 14698 | 2387 | 4354 | 12464 |
| Adenosine | C10H13N5O4 | 206212752 | 77492224 | 83812056 | 41731016 | 48310792 | 59866204 | 71605448 | 64222896 |
| Sarcosine | C3H7NO2 | 12490895 | 6819082 | 16798760 | 14030295 | 14413499 | 25982224 | 14779661 | 14054683 |
| Guanosine monophosphate | C10H14N5O8P | 2580694 | 3854853 | 2325076 | 2824962 | 2510093 | 2772497 | 3684708 | 3177060 |
| Citric acid | C6H8O7 | 40533032 | 43773748 | 13771672 | 12141973 | 12900922 | 11108321 | 17249932 | 17997582 |
| 3-Hydroxybutyric acid | C4H8O3 | 1613580 | 1590982 | 1114076 | 700206 | 772588 | 966534 | 7209194 | 900263 |
| Adenosine triphosphate | C10H16N5O13P3 | 86725 | 326480 | 32371 | 14707 | 59256 | 12661 | 17212 | 51922 |
| Guanosine diphosphate | C10H15N5O11P2 | 81393 | 203347 | 45080 | 34507 | 74665 | 30022 | 38104 | 64109 |
| Xanthylic acid | C10H13N4O9P | 16574 | 36603 | 4495 | 5192 | 24352 | 3657 | 4967 | 6174 |
| L-Glutamine | C5H10N2O3 | 590475264 | 592258752 | 329306656 | 644282240 | 427255168 | 603582272 | 457034080 | 344788800 |
| 4,5-Dihydroorotic acid | C5H6N2O4 | 63397 | 24833 | 182904 | 531722 | 415707 | 95802 | 122824 | 202437 |
| Ureidosuccinic acid | C5H8N2O5 | 81726 | 168058 | 222510 | 546900 | 435327 | 82609 | 41401 | 331226 |
| dTDP | C10H16N2O11P2 | 8886 | 11847 | 0 | 0 | 0 | 0 | 0 | 10025 |
| L-Cystine | C6H12N2O4S2 | 40615 | 30416 | 42005 | 31829 | 35818 | 39846 | 43694 | 35929 |
| Cytidine triphosphate | C9H16N3O14P3 | 3931 | 11041 | 1333 | 0 | 0 | 0 | 0 | 0 |

| | | | | | | | | | |
|---|---|---|---|---|---|---|---|---|---|
| Acetyl-CoA | C23H38N7O17P3S | 0 | 0 | 12350 | 0 | 0 | 0 | 0 | 0 |
| NADH | C21H29N7O14P2 | 7399 | 28225 | 10468 | 0 | 4458 | 0 | 0 | 0 |
| NADP | C21H29N7O17P3 | 14371 | 36576 | 13675 | 8363 | 16076 | 8404 | 12396 | 14377 |
| Palmitic acid | C16H32O2 | 1487343 | 8610343 | 7686168 | 2452912 | 1486516 | 7632358 | 1358540 | 6912370 |
| Uridine triphosphate | C9H15N2O15P3 | 5449 | 16604 | 2347 | 0 | 2966 | 0 | 0 | 1866 |
| 5-Phosphoribosylamine | C5H12NO7P | 7526 | 11000 | 3846 | 3923 | 4289 | 11361 | 9990 | 3840 |
| Oxidized glutathione | C20H32N6O12S2 | 13698561 | 20110978 | 6895237 | 10013955 | 8149196 | 9372823 | 8770878 | 11188553 |
| Pantothenic acid | C9H17NO5 | 12548060 | 21957578 | 18956060 | 14171901 | 9051079 | 26008228 | 7282997 | 18721240 |
| N-Acetylglutamic acid | C7H11NO5 | 241081 | 292623 | 164767 | 162656 | 201655 | 60085 | 156910 | 193095 |
| L-Malic acid | C4H6O5 | 101248808 | 96536672 | 90489560 | 109160136 | 103008856 | 92723744 | 101122136 | 92306384 |
| Orotic acid | C5H4N2O4 | 211178 | 98339 | 597985 | 153385 | 397547 | 276791 | 171908 | 158428 |
| Choline | C5H14NO | 2499586 | 4325058 | 2189704 | 627159 | 1527953 | 2972991 | 1515188 | 1641417 |
| Valerylcarnitine | C12H23NO4 | 3734882 | 4771120 | 4269257 | 3969687 | 2481090 | 6283290 | 16664314 | 3410353 |
| Glycerol 3-phosphate | C3H9O6P | 1324287 | 3059123 | 1286738 | 1126607 | 3131935 | 937920 | 521290 | 972742 |
| Fumaric acid | C4H4O4 | 6157775 | 4723282 | 4953514 | 5009855 | 5541052 | 4862652 | 6244748 | 4707131 |
| Fructose 1,6-bisphosphate | C6H14O12P2 | 83784 | 296216 | 18511 | 19518 | 43357 | 59182 | 98264 | 55614 |
| Uridine diphosphate-N-acetylglucosamine | C17H27N3O17P2 | 443775 | 520526 | 459343 | 482204 | 436748 | 441965 | 452681 | 517833 |
| Inosine | C10H12N4O5 | 36877140 | 29935712 | 28431726 | 38132716 | 34556168 | 40987764 | 40046064 | 19468462 |
| Thymidine | C10H14N2O5 | 52608 | 42912 | 34700 | 33338 | 38770 | 27099 | 38573 | 46253 |
| 4-Hydroxyproline | C5H9NO3 | 20323618 | 13735475 | 36437452 | 39270156 | 20075206 | 24173758 | 12216052 | 10045861 |
| L-Cystathionine | C7H14N2O4S | 2347242 | 3426980 | 87030 | 1580736 | 195558 | 101577 | 1258280 | 1360594 |
| Cytidine monophosphate | C9H14N3O8P | 2783558 | 3302124 | 588732 | 2316775 | 841784 | 255342 | 2290674 | 3861244 |
| Adenine | C5H5N5 | 45701984 | 67392112 | 59229004 | 47093400 | 47237916 | 103067360 | 37963768 | 47062308 |
| Propionylcarnitine | C10H19NO4 | 19642168 | 13380926 | 17526766 | 22028082 | 21194530 | 7787948 | 15820732 | 8047440 |
| Glycine | C2H5NO2 | 19415354 | 20801474 | 31956046 | 26256886 | 29078760 | 24726018 | 26989658 | 21344338 |
| D-Glyceraldehyde 3-phosphate | C3H7O6P | 78940 | 81432 | 112086 | 115230 | 88833 | 93229 | 57503 | 483185 |
| Beta-Alanine | C3H7NO2 | 8657988 | 7597574 | 10100660 | 12285298 | 2518169 | 8594570 | 2557615 | 7460599 |
| FAD | C27H33N9O15P2 | 41325 | 46688 | 31198 | 24006 | 22767 | 44379 | 27450 | 27473 |
| Guanine | C5H5N5O | 367574 | 355622 | 239085 | 136347 | 289045 | 169669 | 255121 | 235261 |
| Hypotaurine | C2H7NO2S | 6960173 | 8494799 | 9904720 | 1939654 | 11167063 | 3499530 | 5124637 | 10023533 |
| Uridine 5'-monophosphate | C9H13N2O9P | 10187008 | 9795128 | 5522381 | 9625104 | 5136038 | 2536355 | 8171701 | 10115595 |
| Dihydroxyacetone phosphate | C3H7O6P | 110074 | 109443 | 113939 | 162438 | 129547 | 66954 | 83861 | 154226 |
| Niacinamide | C6H6N2O | 551668992 | 578452480 | 469549600 | 457364416 | 435707968 | 559465088 | 563188928 | 523435104 |
| L-Serine | C3H7NO3 | 19548364 | 17527106 | 22685694 | 27478642 | 24332978 | 20474694 | 16879788 | 15157439 |
| 1-Methylnicotinamide | C7H8N2O | 3700929 | 3906071 | 923402 | 2255375 | 3464313 | 1481074 | 2388564 | 1120292 |
| Ornithine | C5H12N2O2 | 6839692 | 7460647 | 4148241 | 3259201 | 6928986 | 5343197 | 5066647 | 4743069 |
| Creatinine | C4H7N3O | 60775956 | 63114980 | 57082316 | 56925116 | 36901948 | 54342276 | 35690160 | 38550584 |
| Cytidine | C9H13N3O5 | 13969495 | 11489191 | 3606846 | 9760937 | 4517506 | 7634725 | 12078348 | 7449790 |
| Acetylcysteine | C5H9NO3S | 303637 | 174224 | 322067 | 243746 | 280759 | 240725 | 347281 | 255840 |
| D-Sedoheptulose 7-phosphate | C7H15O10P | 500310 | 1341642 | 427817 | 365302 | 284836 | 506282 | 706414 | 719389 |
| D-Ribose 5-phosphate | C5H11O8P | 678247 | 843468 | 459286 | 318759 | 592284 | 900556 | 889140 | 308667 |
| D-Glucose | C6H12O6 | 1157880 | 962618 | 1055189 | 585503 | 780006 | 808030 | 195882 | 193380 |
| N-Acetyl-L-methionine | C7H13NO3S | 643153 | 916973 | 869445 | 432572 | 925288 | 1341174 | 1321671 | 654505 |
| Carbamoyl phosphate | CH4NO5P | 6643362 | 7512084 | 3752634 | 4352498 | 5895716 | 5194788 | 5871304 | 6031521 |
| Taurine | C2H7NO3S | 213389984 | 239113328 | 171969280 | 174279808 | 138530720 | 192994272 | 175686848 | 199545520 |
| Oxoglutaric acid | C5H6O5 | 281931 | 523071 | 195444 | 147884 | 273170 | 156615 | 338404 | 295802 |
| S-Adenosylmethionine | C15H22N6O5S | 970947 | 1156222 | 2225356 | 1607474 | 3307586 | 2420248 | 1193121 | 1564493 |
| Succinic acid | C4H6O4 | 10753147 | 12995722 | 5751782 | 15049164 | 9761845 | 14511003 | 8548470 | 9137209 |
| Uridine 5'-diphosphate | C9H14N2O12P2 | 81491 | 121872 | 65009 | 65323 | 67794 | 63584 | 73154 | 118472 |
| D-2-Hydroxyglutaric acid | C5H8O5 | 6803682 | 8540664 | 5844392 | 6298418 | 8029147 | 5712135 | 8234748 | 5820260 |
| Adenosine monophosphate | C10H14N5O7P | 67467344 | 81589744 | 57984104 | 78486744 | 61847452 | 58831648 | 60330488 | 78250856 |
| L-Lysine | C6H14N2O2 | 16848564 | 16547048 | 7428940 | 11295214 | 16535060 | 12857676 | 11896642 | 14931688 |
| Adenylsuccinic acid | C14H18N5O11P | 367825 | 546043 | 216321 | 205495 | 170468 | 261506 | 336680 | 313402 |
| Creatine | C4H9N3O2 | 1516729728 | 1368985344 | 1500627200 | 1669640448 | 1474162176 | 1870825088 | 1530600960 | 1341370240 |
| N-Acetyl-L-alanine | C5H9NO3 | 10700887 | 8654781 | 4695366 | 4032745 | 2551257 | 5109394 | 7979453 | 6508177 |
| L-Aspartic acid | C4H7NO4 | 50930956 | 54321752 | 55616348 | 53795100 | 54042400 | 43392356 | 54042888 | 50471028 |
| 2,3-Diphosphoglyceric acid | C3H8O10P2 | 3032 | 2637 | 0 | 0 | 0 | 0 | 0 | 0 |
| Cytosine | C4H5N3O | 0 | 0 | 0 | 27144 | 20505 | 0 | 41530 | 74994 |
| Dihydrofolic acid | C19H21N7O6 | 0 | 0 | 0 | 0 | 0 | 0 | 0 | 0 |
| Folic acid | C19H19N7O6 | 0 | 0 | 0 | 0 | 0 | 0 | 0 | 0 |
| Glutathione | C10H17N3O6S | 31474 | 0 | 0 | 0 | 51566 | 28679 | 0 | 30283 |
| Guanosine triphosphate | C10H16N5O14P3 | 2084 | 8713 | 1271 | 0 | 1294 | 0 | 0 | 1415 |
| Homocysteine | C4H9NO2S | 7869 | 8648 | 8859 | 10136 | 5347 | 3773 | 3945 | 6372 |
| L-Arginine | C6H14N4O2 | 19850676 | 22756586 | 12195267 | 17252066 | 21426932 | 22555348 | 16681076 | 24964050 |
| L-Cysteine | C3H7NO2S | 2919 | 2677 | 3067 | 2803 | 1999 | 2576 | 2891 | 3432 |
| N6-Acetyl-L-lysine | C8H16N2O3 | 58430 | 63961 | 1385730 | 1765371 | 1104993 | 126450 | 2593928 | 1409405 |
| NADPH | C21H30N7O17P3 | 0 | 0 | 0 | 1681 | 0 | 0 | 0 | 0 |
| Nicotinic acid | C6H5NO2 | 27105 | 18564 | 18126 | 17189 | 14339 | 16587 | 13273 | 16849 |
| Orotidylic acid | C10H13N2O11P | 0 | 0 | 0 | 0 | 0 | 0 | 0 | 0 |
| Oxoadipic acid | C6H8O5 | 3235 | 2747 | 3466 | 5459 | 3036 | 3093 | 3144 | 6917 |
| Phenol | C6H6O | 182064 | 254383 | 436496 | 221271 | 228917 | 311313 | 197848 | 414292 |
| Phosphoribosyl pyrophosphate | C5H13O14P3 | 1002 | 0 | 0 | 0 | 0 | 0 | 1065 | 1232 |
| Saccharopine | C11H20N2O6 | 0 | 0 | 0 | 1625 | 0 | 2719 | 0 | 0 |
| Uric acid | C5H4N4O3 | 0 | 0 | 0 | 0 | 0 | 98083 | 0 | 0 |

| KL #3 | ttest_pval | 1/pvalue | KL_Mean | KP_Mean | Log2FoldChange | LogFoldChange | blank_threshold | combined_mean |
|---|---|---|---|---|---|---|---|---|
| 77568952 | 0.00108414 | 922.394097 | 71360730.7 | 115119008 | -0.6899238 | 0.6198866 | 1223953 | 93239869.3 |
| 1233300 | 0.00178452 | 560.375767 | 977291 | 2844318.33 | -1.5412228 | 0.3435941 | 26619 | 1910804.67 |
| 1892761 | 0.00297071 | 336.619862 | 1940498 | 877415.333 | 1.14509512 | 2.21160712 | 35024 | 1408956.67 |
| 3932189 | 0.00858485 | 116.484252 | 3711595 | 10768068 | -1.5366482 | 0.34468532 | 103704 | 7239831.5 |
| 20603828 | 0.00980755 | 101.962236 | 23769191.3 | 57630429.3 | -1.2777379 | 0.41244168 | 1550336.5 | 40699810.3 |
| 126531560 | 0.01308076 | 76.4481713 | 76203021.3 | 187756171 | -1.3009402 | 0.40586161 | 15229640.5 | 131979596 |
| 410140800 | 0.01421246 | 70.3607787 | 337267653 | 556132021 | -0.7215334 | 0.6064525 | 43015967 | 446699837 |
| 106337224 | 0.02850511 | 35.0814324 | 100569555 | 172167637 | -0.7756204 | 0.58413739 | 6078067 | 136368596 |
| 1861214 | 0.03324854 | 30.0765053 | 1479668 | 578691.333 | 1.35440756 | 2.55692096 | 79620 | 1029179.67 |
| 301500096 | 0.03523618 | 28.3799204 | 294872619 | 268340000 | 0.13602974 | 1.09887687 | 20040628 | 281606309 |
| 1918850 | 0.03610551 | 27.696604 | 1386044.67 | 3233965.67 | -1.2223306 | 0.42858979 | 38760.5 | 2310005.17 |
| 73538096 | 0.04582558 | 21.8218723 | 58018896 | 132639171 | -1.1929121 | 0.43741902 | 2779116 | 95329033.3 |
| 3186069 | 0.04680475 | 21.3653552 | 2416862.67 | 4067886.33 | -0.7511439 | 0.5941323 | 630693.5 | 3242374.5 |
| 7974379 | 0.04795415 | 20.8532543 | 5615584.67 | 9345773 | -0.7348778 | 0.60086893 | 1135660.5 | 7480678.83 |
| 1178452480 | 0.05371822 | 18.6156575 | 963330091 | 529310144 | 0.86391693 | 1.81997285 | 21327858 | 746320117 |
| 84591048 | 0.06738061 | 14.8410646 | 92670568 | 154661835 | -0.7389341 | 0.59918187 | 4542025 | 123666201 |
| 30915 | 0.06932173 | 14.4254919 | 25667.3333 | 32676 | -0.348298 | 0.78551026 | 10000 | 29171.6667 |
| 80499800 | 0.07736273 | 12.9261215 | 86475040 | 136021523 | -0.6534793 | 0.63574527 | 3427478 | 111248281 |
| 24112352 | 0.09416405 | 10.619764 | 18751890.3 | 25881484 | -0.4648843 | 0.72452918 | 868886.5 | 22316687.2 |
| 1538931 | 0.10741915 | 9.30932696 | 1973175.33 | 3749318 | -0.926109 | 0.5262758 | 22745 | 2861246.67 |
| 5157989 | 0.11510163 | 8.68797425 | 5299111 | 7558391.67 | -0.5123289 | 0.70108976 | 29392.5 | 6428751.33 |
| 46262 | 0.11965362 | 8.35745745 | 67570 | 228227.333 | -1.7560168 | 0.29606445 | 35981 | 147898.667 |
| 26338268 | 0.12389915 | 8.07108035 | 23016868.7 | 15864796.7 | 0.53686254 | 1.45081397 | 35242 | 19440832.7 |
| 207345216 | 0.12415727 | 8.05430094 | 191683205 | 274971259 | -0.5205569 | 0.69710269 | 4338240 | 233327232 |
| 5270696 | 0.12515191 | 7.99028968 | 4908181.33 | 6808423.33 | -0.4721322 | 0.72089838 | 79913 | 5858302.33 |
| 73678 | 0.13279125 | 7.5306165 | 70815 | 131658 | -0.8946683 | 0.53787085 | 14464.5 | 101236.5 |
| 626501248 | 0.13936667 | 7.17531663 | 566530293 | 868769515 | -0.6168204 | 0.65210655 | 53788525.5 | 717649904 |
| 451165 | 0.14325653 | 6.98048443 | 520903 | 305373.667 | 0.77043909 | 1.70578886 | 126072.5 | 413138.333 |
| 23573 | 0.14674289 | 6.81464037 | 21970 | 30320.3333 | -0.4647507 | 0.72459626 | 12024.5 | 26145.1667 |
| 223053 | 0.15289294 | 6.54052431 | 226310.333 | 158695 | 0.51204579 | 1.42607097 | 31423 | 192502.667 |
| 113339 | 0.15311753 | 6.53093086 | 176432 | 336811.333 | -0.9328284 | 0.52383035 | 37737.5 | 256621.667 |
| 454832 | 0.15734774 | 6.35535038 | 473515 | 873037.667 | -0.8826338 | 0.54237637 | 473515 | 673276.333 |
| 91235368 | 0.15951744 | 6.26890694 | 69132238.7 | 96682517.3 | -0.4838964 | 0.71504384 | 17408392.5 | 82907378 |
| 1514177664 | 0.17271456 | 5.78989996 | 1243218325 | 984515413 | 0.33659398 | 1.26277183 | 14852409.5 | 1113866869 |
| 28746 | 0.17431725 | 5.73666699 | 27342 | 35657.6667 | -0.3830935 | 0.76679162 | 12902 | 31499.8333 |
| 120421 | 0.18183378 | 5.49952815 | 102373.667 | 418723.667 | -2.0321538 | 0.2444898 | 40375 | 260548.667 |
| 388619 | 0.18725468 | 5.3403205 | 368437 | 962393 | -1.3852082 | 0.38283425 | 34830 | 665415 |
| 707734 | 0.19086959 | 5.23917916 | 1574910 | 373127.333 | 2.07752943 | 4.22083793 | 84697 | 974018.667 |
| 333660 | 0.19632275 | 5.09365306 | 468508 | 641220 | -0.4527457 | 0.73065095 | 32625.5 | 554864 |
| 208801 | 0.1981943 | 5.04555378 | 97972 | 1188440.67 | -3.6005566 | 0.08243743 | 18511.5 | 643206.333 |
| 7837002 | 0.2012008 | 4.97015908 | 4892560.33 | 2637407.33 | 0.89146924 | 1.85506435 | 76360.5 | 3764983.83 |
| 12062870 | 0.20123386 | 4.96934263 | 15479077 | 38315038.7 | -1.3075913 | 0.40399482 | 3187147 | 26897057.8 |
| 1015139 | 0.20123508 | 4.96931239 | 1853384.33 | 640888 | 1.53201792 | 2.89190051 | 106726.5 | 1247136.17 |
| 151482656 | 0.207916 | 4.80963459 | 136881995 | 152814203 | -0.1588459 | 0.89574131 | 3804363 | 144848099 |
| 11427035 | 0.20819786 | 4.80312344 | 15654842 | 17129536.7 | -0.1298772 | 0.91390925 | 15654842 | 16392189.3 |
| 812040000 | 0.20840903 | 4.79825667 | 734891136 | 534999552 | 0.45799287 | 1.37362944 | 5889004.5 | 634945344 |
| 23193876 | 0.20851069 | 4.79591714 | 23292694.7 | 30239659.3 | -0.3765643 | 0.77026974 | 5877015 | 26766177 |
| 7870056 | 0.21142198 | 4.72987718 | 7714020 | 11267343 | -0.5465926 | 0.68463523 | 141260.5 | 9490681.5 |
| 1376140 | 0.21194616 | 4.71817941 | 1700114.67 | 1308288.67 | 0.37795116 | 1.29949507 | 31758.5 | 1504201.67 |
| 140459 | 0.22312145 | 4.48186398 | 83946.6667 | 252484 | -1.588647 | 0.33248311 | 31732 | 168215.333 |
| 9629 | 0.22348352 | 4.47460279 | 15499 | 29285.3333 | -0.9180032 | 0.52924103 | 10000 | 22392.1667 |
| 3231139 | 0.23023824 | 4.34332721 | 2862790.33 | 3724741.67 | -0.3797184 | 0.76858762 | 190166 | 3293766 |
| 607883 | 0.24622357 | 4.06134964 | 849205 | 2322786.67 | -1.4516719 | 0.3655975 | 443622 | 1585995.83 |
| 10004815 | 0.24631471 | 4.05984686 | 9124193.33 | 10544087 | -0.2086653 | 0.86533745 | 47770.5 | 9834140.17 |
| 17389676 | 0.25185901 | 3.97047545 | 14210817.3 | 19343918.3 | -0.4448905 | 0.73464006 | 1588867 | 16777367.8 |
| 22363 | 0.25190948 | 3.96967983 | 64237.8333 | 35368.5 | 0.8609582 | 1.81624421 | 22957.5 | 49803.1667 |
| 4603138 | 0.25235238 | 3.9627127 | 2642474 | 1053767.67 | 1.32633246 | 2.50764384 | 35368.5 | 1848120.83 |
| 1340374 | 0.25903859 | 3.86042865 | 1893801.67 | 1278428.67 | 0.56691358 | 1.48135106 | 117834 | 1586115.17 |
| 7067 | 0.26150223 | 3.82405919 | 19363 | 29863.6667 | -0.6250888 | 0.64837986 | 19363 | 24613.3333 |
| 67983256 | 0.26317925 | 3.7996917 | 67937200 | 122505677 | -0.8505749 | 0.55456369 | 617523 | 95221438.7 |
| 25138036 | 0.26523146 | 3.77029176 | 17990793.3 | 12036245.7 | 0.57987335 | 1.49471802 | 2661445.5 | 15013519.5 |
| 3792916 | 0.28325081 | 3.53044006 | 3551561.33 | 2920207.67 | 0.28238243 | 1.21620163 | 225652.5 | 3235884.5 |
| 26099780 | 0.28466762 | 3.51286874 | 20449098 | 32692817.3 | -0.6769365 | 0.62549207 | 4528381 | 26570957.7 |
| 2988877 | 0.29178098 | 3.42722818 | 3699444.67 | 1439546 | 1.36169483 | 2.56986902 | 153075.5 | 2569495.33 |
| 25161 | 0.29285651 | 3.41464158 | 39174 | 148668.333 | -1.924129 | 0.26349929 | 32800 | 93921.1667 |
| 41334 | 0.29559868 | 3.38296499 | 59897.6667 | 114177.333 | -0.9307046 | 0.52460208 | 57792 | 87037.5 |
| 15355 | 0.32112748 | 3.11402812 | 11785 | 21059 | -0.8374852 | 0.55961822 | 10000 | 16422 |
| 396508032 | 0.32458204 | 3.08088516 | 399443637 | 504013557 | -0.3354706 | 0.79252558 | 71465331 | 451728597 |
| 3735277 | 0.34905767 | 2.86485609 | 1353512.67 | 90378 | 3.90459295 | 14.9761299 | 12537.5 | 721945.333 |
| 4686920 | 0.36654094 | 2.72820818 | 1686515.67 | 157431.333 | 3.42125111 | 10.7127065 | 13121.5 | 921973.5 |
| 0 | 0.37390097 | 2.67450499 | 11727.5 | 11767.3333 | -0.0048919 | 0.99661492 | 11727.5 | 11747.4167 |
| 88710 | 0.37390097 | 2.67450499 | 87495.6667 | 86888.5 | 0.01004632 | 1.00698788 | 86888.5 | 87192.0833 |
| 0 | 0.37390097 | 2.67450499 | 10578 | 10732.3333 | -0.0208969 | 0.98561978 | 10578 | 10655.1667 |

| | | | | | | | | |
|---|---|---|---|---|---|---|---|---|
| 0 | 0.37390097 | 2.67450499 | 10000 | 10783.3333 | -0.1088032 | 0.92735703 | 10000 | 10391.6667 |
| 0 | 0.37390097 | 2.67450499 | 10778 | 16593.6667 | -0.6225432 | 0.64952492 | 10778 | 13685.8333 |
| 16587 | 0.37390097 | 2.67450499 | 19700.5 | 25325.6667 | -0.362368 | 0.77788673 | 19700.5 | 22513.0833 |
| 17109180 | 0.37390097 | 2.67450499 | 14857853.7 | 13732190.5 | 0.11366395 | 1.08197259 | 13732190.5 | 14295022.1 |
| 0 | 0.37390097 | 2.67450499 | 11011.5 | 12875.6667 | -0.2256361 | 0.85521785 | 11011.5 | 11943.5833 |
| 4877 | 0.37390097 | 2.67450499 | 10579.5 | 10719.6667 | -0.0189886 | 0.98692434 | 10579.5 | 10649.5833 |
| 9431970 | 0.38647263 | 2.58750537 | 9797133.67 | 13568258.7 | -0.469804 | 0.72206271 | 536280 | 11682696.2 |
| 14791851 | 0.38715027 | 2.5829764 | 13598696 | 17820566 | -0.3900748 | 0.76309002 | 27258.5 | 15709631 |
| 226404 | 0.38970572 | 2.56603882 | 192136.333 | 232823.667 | -0.2771074 | 0.825244 | 24057.5 | 212480 |
| 115635496 | 0.40666715 | 2.45901349 | 103021339 | 96091680 | 0.10045976 | 1.07211507 | 9590830 | 99556509.3 |
| 5315364 | 0.41128506 | 2.43140368 | 1881900 | 302500.667 | 2.63717974 | 6.22114331 | 12018.5 | 1092200.33 |
| 3388002 | 0.41182854 | 2.42819499 | 2181535.67 | 3004782.67 | -0.4619166 | 0.72602112 | 484320.5 | 2593159.17 |
| 4363533 | 0.41492968 | 2.41004691 | 8146066.67 | 4258419.67 | 0.93578549 | 1.91293186 | 21861 | 6202243.17 |
| 2125868 | 0.41656118 | 2.40060773 | 1206633.33 | 1890049.33 | -0.6474365 | 0.63841367 | 162414 | 1548341.33 |
| 7174355 | 0.41751349 | 2.39513219 | 6042078 | 5278190.33 | 0.19500143 | 1.1447253 | 242306.5 | 5660134.17 |
| 41260 | 0.42868814 | 2.33269805 | 65046 | 137591.333 | -1.0808574 | 0.4727478 | 32774 | 101318.667 |
| 547892 | 0.4365947 | 2.29045382 | 506135.333 | 474548 | 0.09296917 | 1.06656299 | 22253 | 490341.667 |
| 15054997 | 0.44426764 | 2.25089541 | 24856507.7 | 31748192.7 | -0.3530509 | 0.7829267 | 678816 | 28302350.2 |
| 81035 | 0.44544595 | 2.24494129 | 55287 | 43406.6667 | 0.34902365 | 1.27369836 | 18619 | 49346.8333 |
| 26829814 | 0.45134904 | 2.21558019 | 16363909 | 23498848.3 | -0.5220726 | 0.69637068 | 6147403 | 19931378.7 |
| 743976 | 0.45277708 | 2.20859234 | 1120950 | 1953750.67 | -0.8015244 | 0.57374261 | 28492 | 1537350.33 |
| 2846684 | 0.46067027 | 2.17075003 | 2999534 | 2224804.67 | 0.43105971 | 1.34822353 | 54164.5 | 2612169.33 |
| 64019984 | 0.47760313 | 2.09378864 | 49682020 | 57441033.3 | -0.2093579 | 0.86492211 | 488887.5 | 53561526.7 |
| 18366572 | 0.48514935 | 2.06122094 | 14078248 | 16849953.3 | -0.2592768 | 0.83550665 | 50544.5 | 15464100.7 |
| 38496588 | 0.48907968 | 2.04465663 | 28943528 | 24057624.7 | 0.26674658 | 1.20309168 | 3702793.5 | 26500576.3 |
| 52907 | 0.49591129 | 2.01648969 | 197865 | 90819.3333 | 1.12344509 | 2.17866607 | 23931 | 144342.167 |
| 10797044 | 0.50108981 | 1.99565026 | 6938419.33 | 8785407.33 | -0.3405021 | 0.78976638 | 229027 | 7861913.33 |
| 47378 | 0.52207126 | 1.91544733 | 34100.3333 | 39737 | -0.2206971 | 0.85815067 | 10000 | 36918.6667 |
| 355843 | 0.52370108 | 1.90948621 | 282075 | 320760.333 | -0.1854169 | 0.8793949 | 43114.5 | 301417.667 |
| 18438098 | 0.52866846 | 1.89154466 | 11195422.7 | 8453230.67 | 0.40533427 | 1.32439574 | 209699 | 9824326.67 |
| 9853556 | 0.61488291 | 1.62632589 | 9380284 | 8501505.67 | 0.14191323 | 1.10336738 | 294481.5 | 8940894.83 |
| 47481 | 0.63756073 | 1.56847803 | 95189.3333 | 111152 | -0.2236621 | 0.85638885 | 34610 | 103170.667 |
| 566733504 | 0.6414322 | 1.55901123 | 551119179 | 533223691 | 0.04762345 | 1.03356094 | 2248355.5 | 542171435 |
| 23514488 | 0.65985197 | 1.51549143 | 18517238.3 | 19920388 | -0.1053768 | 0.92956213 | 2445214 | 19218813.2 |
| 3403211 | 0.66785757 | 1.49732524 | 2304022.33 | 2843467.33 | -0.3034965 | 0.8102862 | 813497.5 | 2573744.83 |
| 7409998 | 0.70247237 | 1.42354354 | 6634404.67 | 6848982.33 | -0.0459226 | 0.96867014 | 6246608 | 6741693.5 |
| 149306144 | 0.72395949 | 1.38129276 | 74515629.3 | 60324417.3 | 0.30480098 | 1.23524822 | 2479342 | 67420023.3 |
| 13584517 | 0.72890983 | 1.37191181 | 11037551.7 | 9688510.67 | 0.18807338 | 1.13924132 | 63430.5 | 10363031.2 |
| 250229 | 0.76701393 | 1.30375728 | 284450 | 266642.667 | 0.09326743 | 1.06678351 | 11558 | 275546.333 |
| 607754 | 0.80298608 | 1.24535161 | 677852.333 | 756589.667 | -0.15854 | 0.89593126 | 60596 | 717221 |
| 629710 | 0.81200284 | 1.23152279 | 609172.333 | 660333.667 | -0.1163448 | 0.922522 | 45313.5 | 634753 |
| 3570552 | 0.82793889 | 1.2078186 | 1319938 | 1058562.33 | 0.31836394 | 1.24691571 | 33160.5 | 1189250.17 |
| 622093 | 0.82844122 | 1.20708624 | 866089.667 | 809857 | 0.09684921 | 1.0694353 | 10873.5 | 837973.333 |
| 6692536 | 0.84395549 | 1.18489661 | 6198453.67 | 6365539.83 | -0.0383745 | 0.97375145 | 4941173.5 | 6281996.75 |
| 233633568 | 0.8499984 | 1.1764728 | 202955312 | 208157531 | -0.0365137 | 0.97500826 | 7369847.5 | 205556421 |
| 412313 | 0.88950358 | 1.12422257 | 348839.667 | 333482 | 0.06495521 | 1.04605246 | 25355.5 | 341160.833 |
| 1438101 | 0.90373826 | 1.10651507 | 1398571.67 | 1450841.67 | -0.0529359 | 0.96397264 | 318604 | 1424706.67 |
| 11044215 | 0.91533033 | 1.09250176 | 9576631.33 | 9833550.33 | -0.0381941 | 0.97387322 | 517644 | 9705090.83 |
| 83585 | 0.92157105 | 1.08510353 | 91737 | 89457.3333 | 0.03630398 | 1.02548328 | 15545 | 90597.1667 |
| 6816308 | 0.92495105 | 1.08113829 | 6957105.33 | 7062912.67 | -0.0217761 | 0.98501931 | 131471 | 7010009 |
| 66287772 | 0.93730401 | 1.06688971 | 68289705.3 | 69013730.7 | -0.0152153 | 0.98950897 | 6058723.5 | 68651718 |
| 17819660 | 0.96957112 | 1.03138386 | 14882663.3 | 14981069.2 | -0.0095079 | 0.99343132 | 11547595.5 | 14931866.3 |
| 493190 | 0.97047764 | 1.03042044 | 381090.667 | 376729.667 | 0.01660463 | 1.01157594 | 63729 | 378910.167 |
| 1523377408 | 0.97099548 | 1.02987091 | 1465116203 | 1462114091 | 0.0029592 | 1.00205327 | 354029448 | 1463615147 |
| 9451063 | 0.98564819 | 1.01456078 | 7979564.33 | 8017011.33 | -0.0067545 | 0.99532906 | 75022 | 7998287.83 |
| 56237304 | 0.98651962 | 1.01366459 | 53583740 | 53623018.7 | -0.0010572 | 0.9992675 | 2591066 | 53603379.3 |
| 0 | NA | NA | 10000 | 10000 | 0 | 1 | 10000 | 10000 |
| 17975 | NA | NA | 75611 | 75611 | 0 | 1 | 75611 | 75611 |
| 0 | NA | NA | 10000 | 10000 | 0 | 1 | 10000 | 10000 |
| 0 | NA | NA | 10000 | 10000 | 0 | 1 | 10000 | 10000 |
| 0 | NA | NA | 66110 | 66110 | 0 | 1 | 66110 | 66110 |
| 1118 | NA | NA | 10699.5 | 10699.5 | 0 | 1 | 10699.5 | 10699.5 |
| 3818 | NA | NA | 10858 | 10858 | 0 | 1 | 10858 | 10858 |
| 23667016 | NA | NA | 27679190.5 | 27679190.5 | 0 | 1 | 27679190.5 | 27679190.5 |
| 2633 | NA | NA | 18144 | 18144 | 0 | 1 | 18144 | 18144 |
| 5702810 | NA | NA | 10030944.5 | 10030944.5 | 0 | 1 | 10030944.5 | 10030944.5 |
| 2566 | NA | NA | 10000 | 10000 | 0 | 1 | 10000 | 10000 |
| 21462 | NA | NA | 52132 | 52132 | 0 | 1 | 52132 | 52132 |
| 0 | NA | NA | 37149 | 37149 | 0 | 1 | 37149 | 37149 |
| 12038 | NA | NA | 16503.5 | 16503.5 | 0 | 1 | 16503.5 | 16503.5 |
| 271055 | NA | NA | 4626068.5 | 4626068.5 | 0 | 1 | 4626068.5 | 4626068.5 |
| 0 | NA | NA | 10000 | 10000 | 0 | 1 | 10000 | 10000 |
| 3505 | NA | NA | 10470 | 10470 | 0 | 1 | 10470 | 10470 |
| 0 | NA | NA | 10591 | 10591 | 0 | 1 | 10591 | 10591 |
